# Strand- and replication timing-dependent functions of DNA polymerase η in human DNA replication and mutagenesis

**DOI:** 10.1101/2025.09.10.675280

**Authors:** Lewis J. Bainbridge, Yuji Masuda, Kanae Kaneko, Tamiko Minamisawa, Mami Takahashi, Chikahide Masutani, Yasukazu Daigaku

## Abstract

Efficient eukaryotic DNA replication relies on the coordinated actions of replicative and error-prone polymerases, with the latter providing flexibility at the cost of mutagenesis. DNA polymerase η (Pol η) is most recognised for tolerating UV-induced DNA damage via translesion synthesis (TLS). However, emerging evidence suggests a broader contribution of error-prone polymerases to DNA synthesis. To elucidate the roles of Pol η during unperturbed replication, we applied polymerase usage sequencing (Pu-seq) to map its activity genome-wide. Our findings demonstrate that Pol η preferentially participates in lagging-strand replication, consistent with observations in budding yeast. Furthermore, Pol η usage varied throughout S phase, with a pronounced enrichment in late-replicating domains, dependent on PCNA ubiquitylation. Under moderate doses of UV, Pol η usage retained its replicative strand bias, which contrasted with the prominent strand bias observed in transcribed regions resulting from asymmetric repair processes. These results reveal that Pol η’s flexibility and intrinsic coupling with replication forks extend beyond TLS in human cells. In cancer genomes, characteristic Pol η mutations are enriched in late-replicating regions and correlate with RAD18 expression, implicating PCNA-mediated Pol η activation in mutagenesis. Together, these findings reveal an unexpected bias in Pol η usage during unperturbed replication which may represent a key contribution to the mutational burden in the human genome.

## Introduction

Genomic DNA is copied by DNA polymerase enzymes. Cells possess myriad polymerases that coordinate to ensure timely and accurate duplication of the genetic material. In eukaryotes, bulk replication progresses as a replication fork structure, where Pol ε synthesises the continuous leading strand while Pol α and Pol δ coordinate Okazaki fragment synthesis on the lagging strand (reviewed in ^1^). Other polymerases play more specialised roles, offering flexibility to tolerate complications that may arise during replication, such as DNA damage, and are therefore crucial mediators of genomic stability. The most well-studied of these is Pol η, the gene which is deficient in Xeroderma Pigmentosum Variant (XPV) patients^2^. Pol η was primarily identified as the enzyme bypassing UV-induced damage, particularly cyclobutane pyrimidine dimers (CPD), with high accuracy via translesion synthesis (TLS)^3, 4, 5^. Later studies showed that Pol η’s enlarged active site also enables accurate TLS over cisplatin-induced lesions^6, 7^ ^8^. Pol η is also the key DNA polymerase responsible for A/T mutations during somatic hypermutation in B cells, acting downstream of cytosine deaminase–induced mismatch repair^9, 10^. Interestingly, Pol η mRNA is comparably expressed across multiple tissues^11^, not solely in sun-exposed skin or immune cells (**Supplementary** Fig. 1a), suggesting functions in replication beyond TLS. Previous observations support this, with Pol η and travelling with the replication fork^12^ and forming foci in unperturbed cells^13^. Indeed, additional roles of Pol η have been identified, for instance, Pol η is required to maintain common fragile site (CFS) stability, even in the absence of exogenous stress^14^. These structure-forming repeat sequences are inefficiently replicated by Pol δ, but well tolerated by Pol η, which is recruited to these sites^15, 16^. The use of Pol η at these regions of non-B DNA reduces fork pausing^17^. These roles highlight the versatility of Pol η during DNA synthesis, a property that may also help cells tolerate replication stress associated with oncogenic transformation (reviewed in ^18^).

Given the diverse functions of Pol η and other error-prone polymerases, elucidating their roles in genome replication and how they coordinate with the replicative polymerases remains a key focus. To examine polymerase coordination during human replication, we recently established the polymerase usage sequencing (Pu-seq) assay in HCT116 cells^19^. Analysis of Pol ε and Pol α usage in this system revealed unexpected deviations from canonical patterns in specific regions, influenced by transcribed genes and large-scale chromosomal domains^19^. These observations hint towards the involvement of other polymerases, which were not included in the analyses. To date, global analysis of error-prone polymerase usage in eukaryotes are limited to budding yeast Pol η, which was determined to contribute to lagging-strand synthesis^20^. However, the yeast genome is ∼300-fold smaller than that of humans and lacks distinct chromosomal domains. Applying Pu-seq in human cells therefore provides an opportunity to determine roles of polymerases that may be overlooked in the yeast genome. Originally developed in fission yeast^21, 22^, Pu-seq maps polymerase activity by detecting ribonucleotides (rNMPs or rNTP) incorporated into nascent DNA by mutant polymerases with reduced nucleotide selectivity. To prevent removal of these rNMPs, human Pu-seq assays typically employ the auxin-inducible degron 2 (AID2) degron system to degrade RNASEH2A, thereby inactivating the ribonucleotide excision repair (RER) pathway during the experiment^23^. In principle, Pu-seq can be applied to any polymerase, provided a mutant can be generated that incorporates rNMPs at a higher rate than the wild-type (WT) protein. Therefore, we set out to establish a Pol η Pu-seq system to map Pol η synthesis events and investigate its role during human DNA replication.

## Results

### The Pol η-F18A mutation increases ribonucleotide incorporation

To map polymerase usage with Pu-seq, a polymerase mutant with an increased propensity to incorporate rNMPs is required. The specificity of polymerases for deoxy-ribonucleotides (dNTPs) over the more abundant rNTPs is primarily governed by a steric gate residue (tyrosine or phenylalanine in B and Y family polymerases), which sterically excludes the occupancy of nucleotides with a 2′-hydroxyl moiety^24^. In budding yeast Pol η, strict rNTP discrimination is predominantly governed by F35 and substituting this residue for alanine (Pol η-F35A) dramatically increases rNTP incorporation in vitro^25^, which has been used to map Pol η usage in yeast^20^. Although human Pol η is less selective than its yeast counterpart^26^, the steric gate residue (F18) demonstrates high sequence and structural conservation with yeast (**Fig. 1a, Supplementary** Fig. 1b). Structural models indicate that steric hindrance between this residue and an incoming rNTP distances the α-phosphate from the nascent strand terminus, rendering phosphodiester bond formation inefficient^26^. We therefore selected the Pol η-F18A mutation as a candidate for use in Pu-seq assays to attempt to define how Pol η is utilized in human cells.

**Figure 1.**
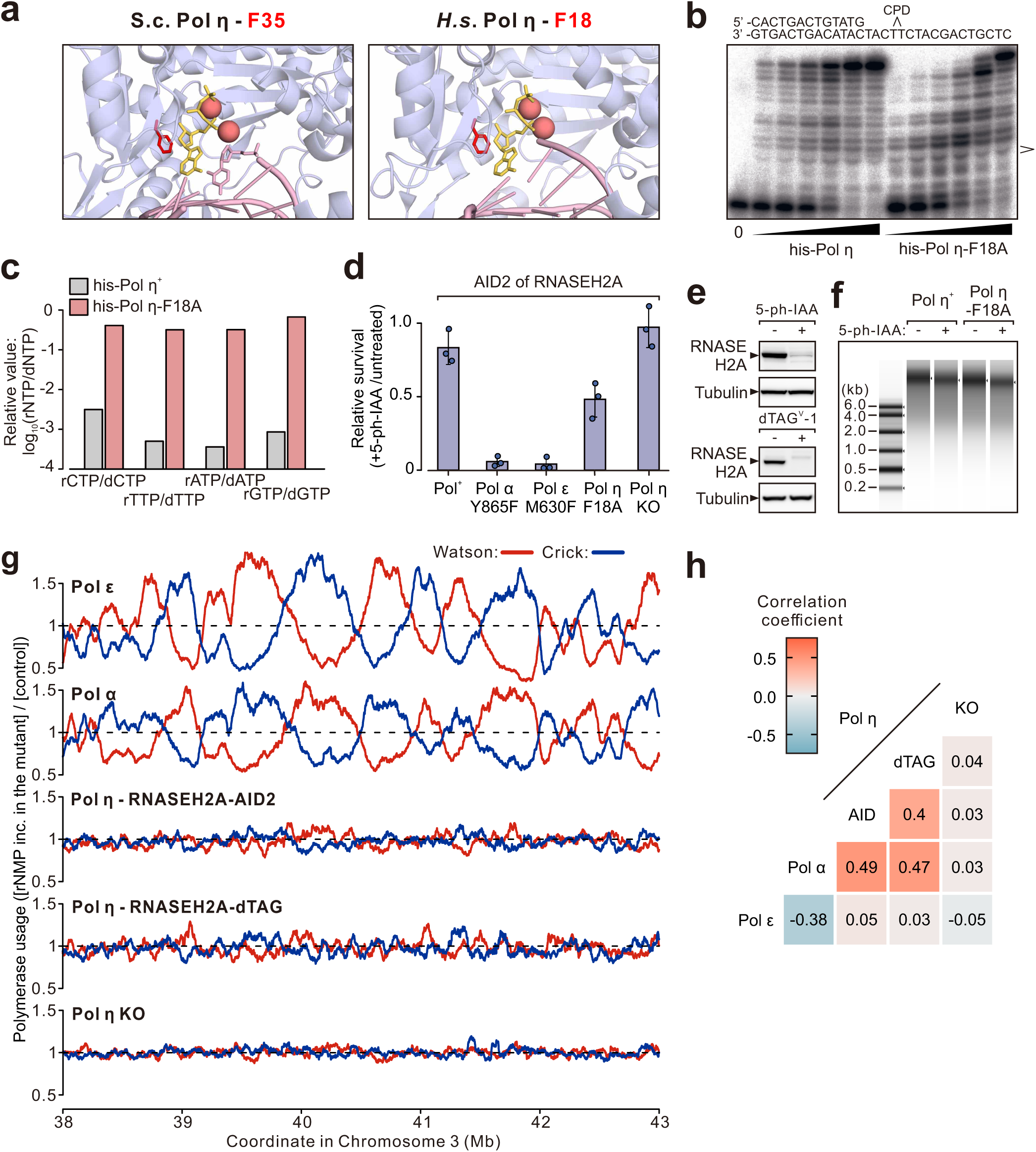
The Pol η usage profile displays a lagging strand bias. **a** Structural comparison of *Saccharomyces ceravisiae* (PDB: 3MFH) and *Homo sapien* (PDB: 3MR2) Pol η active sites. Conserved steric gate phenylalanine residues (red) sterically hinder the 2′-hydroxyl group of incoming nucleotides (yellow sticks). Divalent metal ions (red spheres) and the primed DNA template (pink) are displayed. **b** Polymerase activity of Pol η-F18A over a CPD lesion. A γ-32P-labelled 13-mer primer annealed to a 30-mer template containing a cis-syn thymidine dimer was extended by recombinant His-tagged Pol η or Pol η-F18A in the presence of dNTPs and MgCl_2_. **c** Relative efficiency of rNTP versus dNTP incorporation by Pol η and Pol η-F18A. Relative efficiency was calculated as [*k*_cat_/*K*_m_ rNTP]/[*k*_cat_/*K*_m_ dNTP] for each base, as determined by single-nucleotide incorporation assays. **d** Sensitivity of Pu-seq cells to 7 days of 5-ph-IAA treatment. Survival was assessed by crystal violet staining quantification and normalised to untreated controls. **e** Representative western blots confirming RNaseH2A-mAC degradation after 48-hour 5-ph-IAA treatment (top) and RNaseH2A-dTAG degradation after 48-hour dTAG-^V^1 treatment (bottom). **f** DNA fragmentation following NaOH hydrolysis of DNA harvested from Pu-seq cells after 48 hours of 5-ph-IAA treatment. **g** Strand-specific polymerase usage of Pol ε, Pol α and Pol η across a representative region of Chromosome 3. Red: Watson strand. Blue: Crick strand. Data were smoothed with a moving average (*m* = 30; see Materials and Methods). **h** Heatmap of Pearson correlations between genome-wide strand-specific profiles of Pol ε, Pol α and Pol η variants. Red: positive. Blue: negative.

To evaluate the effect of the F18A mutation, recombinant Pol η was expressed and purified from *Escherichia coli* for in vitro characterisation (**Supplementary** Fig. 2a). Human Pol η-F18A retained polymerase activity with slightly reduced processivity compared to the wild-type enzyme, while performing TLS across a CPD lesion with similar efficiency (**Fig. 1b**). To assess rNTP incorporation, the steady state kinetic parameters for single-nucleotide incorporation were determined (**Table 1, Supplementary** Fig. 2b). Calculation of catalytic efficiency (*k_cat_*/*K_m_*) revealed a striking difference in nucleotide differentiation between the WT and mutant enzyme. The WT enzyme exhibited much higher efficiency for incorporating dNTPs compared to rNTPs. In contrast, the Pol η-F18A mutant exhibited much less distinct preference, with altered kinetic parameters for both dNTPs and rNTPs. Specifically, the mutant enzyme exhibited an increase in *K_m_* for dNTPs and a decrease in *K_m_* for rNTPs, accompanied by an elevation in *k_cat_* for rNTPs, dramatically enhancing the relative efficiency of rNTP incorporation compared to the WT enzyme (**Fig.1c**). These results demonstrate that the Pol η-F18A mutation alters nucleotide incorporation dynamics, which could serve as a detectable signature of Pol η activity in human cells.

**Table 1.**
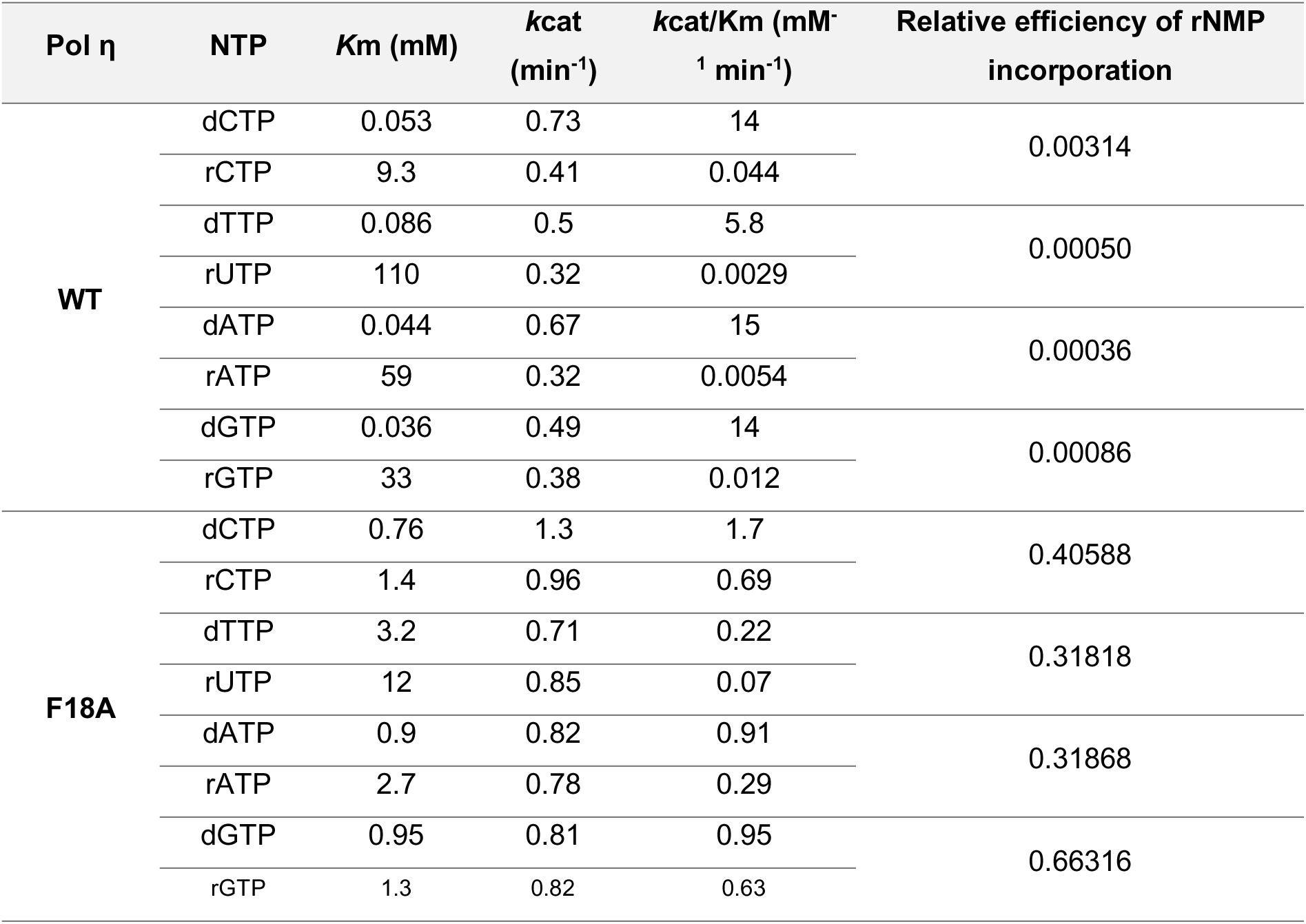
Steady-state kinetic parameters for one-base incorporation by WT and F18A Pol η.

### Establishing Pol η-F18A cell lines

Having established that Pol η-F18A incorporates rNTPs more efficiently than the WT protein, we introduced this mutation into both alleles of the endogenous Pol η-encoding gene (*POLH*) in RNASEH2A-degron HCT116 cells using CRISPR-Cas9 (**Supplementary** Fig. 3a). The mutation did not affect Pol η protein levels (**Supplementary** Fig. 3b) or cell proliferation rate (**Supplementary** Fig. 3c). Pol η-F18A cells also exhibit a cell cycle profile similar to WT, even in the presence of 5-ph-IAA, an auxin compound that induces degradation of RNaseH2A subunit via the AID2 system (**Supplementary** Fig. 3c,d).

To assess functionality of the polymerase mutation in cells, we examined cell fitness under conditions promoting chronic persistence of rNMPs in the genome. We previously demonstrated short-term (∼48 hr) depletion of RNASEH2A via the AID2 system had no significant impact on cell growth, even in cells harbouring rNMP-incorporating replicative polymerases (Pol ε-M630F and Pol α-Y865F). In contrast, long-term (7 days) 5-ph-IAA exposure decreased survival of RNASEH2A-degron cells by ∼20% (**Fig. 1d**). The effect was more severe in Polα-Y865F or Polε-M630F cells, which lost viability, consistent with a high rNMP incorporation-rate of the mutated replicative polymerases. Pol η-F18A cells also exhibited reduced viability, though to a lesser extent, which was not observed in Pol η knockout (KO) cells (**Supplementary** Fig. 3b), reflecting moderate rNTP incorporation by this polymerase.

Given the role of Pol η in DNA damage tolerance, we next assessed cell survival following exposure to UV. Despite the TLS proficiency of Pol η-F18A, Pol η-F18A cells displayed extreme UV sensitivity, exceeding that of Pol η-KO cells (**Supplementary** Fig. 3e). These findings mirror results in budding yeast^25^, suggesting rNMP incorporation opposite or near UV lesions is deleterious for cell survival.

### Detection of genome-wide Pol η usage via Pu-seq

For Pu-seq assays, cells were harvested 48 hours after degrading RNASEH2A to levels barely detectable by western blot (**Fig. 1e**) and genomic DNA was extracted. Genomic DNA was alkaline-treated to hydrolyse rNMPs, and single-stranded DNA was separated by electrophoresis. Unlike replicative polymerase mutants, which produce a marked increase in small DNA fragments^19^, the electrophoresis patterns for Pol η-F18A and control (Pol^+^) samples showed no obvious differences (**Fig. 1f**). This indicates low levels of rNMP incorporation, consistent with the modest viability defect observed during RNASEH2A depletion (**Fig. 1d**). For Pu-seq, alkaline-induced small DNA fragments (<2kb) were isolated to produce libraries for Illumina sequencing. Approximately 200 million paired-end reads were obtained for control and Pol η-F18A cell lines, with the strand-specific 5ʹ end positions denoting rNMP positions, which were scored in 1-kb bins across the genome.

Two independent Pu-seq experiments were conducted with Pol η. The ratio of rNMPs detected in Polη-F18A cells compared to control cells (Pol^+^ with RNASEH2A degradation) was plotted for the Watson and Crick strands along a region of Chromosome 3 (**Fig. 1g, Supplementary** Fig. 4a). As predicted, the Pol η signal intensity was lower than previously observed for Pol ε and Pol α, which contribute to leading and lagging strand synthesis, respectively^19^. However, both replicates showed reproducible fluctuations which were absent in the Pol η-KO control (**Fig. 1g**). To increase the sensitivity of the assay for detecting infrequent polymerase usage, we further decreased RNASEH2A levels by employing the VHL-recruiting dTAG system^27^, achieving degradation to levels undetectable by western blot (**Fig. 1e**). Conducting Pu-seq with this system produced results similar to those obtained with the AID2 system (**Fig. 1g, Supplementary** Fig. 4a), suggesting that the relatively weak Pol η signal reflects its limited usage rather than residual RNASEH2A activity.

### Pol η usage is bias towards the lagging strand

One notable observation was that the Pol η signal visually peaks and troughs in a similar pattern to lagging strand polymerase Pol α. Consistently, strand-specific Pol η scores positively correlate with Pol α genome-wide, but not with leading strand polymerase Pol ε (**Fig. 1h, Supplementary** Fig. 4b). We quantified the strand bias of the Pol η signal by defining a parameter that captures differential polymerase usage (Pu) between the Watson (W) and Crick (C) strands at each genomic position, using a formula analogous to fork directionality: (Pu^W^-Pu^C^)/(Pu^W^+Pu^C^) (**Fig. 2a, Supplementary** Fig. 5)^19, 28, 29^. By examining the pattern of this bias around initiation zones (IZs), where bidirectional replication forks are initiated, we can assess biased assignment of polymerases towards a specific replicative strand (**Fig. 2a**). Consistent with the role of Pol ε in leading strand synthesis and Pol α and lagging strand synthesis, Pol α shows a Watson bias to the left and a Crick bias to the right of all IZs previously identified with Pu-seq, whereas Pol ε shows the opposite pattern (**Fig. 2b, Supplementary** Fig. 6a). Noticeably, Pol η strand bias generally switches from Watson to Crick at these sites, resembling the lagging-strand pattern of Pol α (**Fig. 2b**). The similarity of the Pol η and Pol α profiles is also reflected in the genome-wide correlations (**Supplementary** Fig. 6b). No discernible pattern was observed around IZs in the Pol η-KO control (**Fig. 2b**). Together, these data demonstrate that Pol η usage is biased towards the lagging strand in human cells, consistent with findings in budding yeast^20^, suggesting that Pol η activity is coordinated with replication fork polarity.

**Figure 2.**
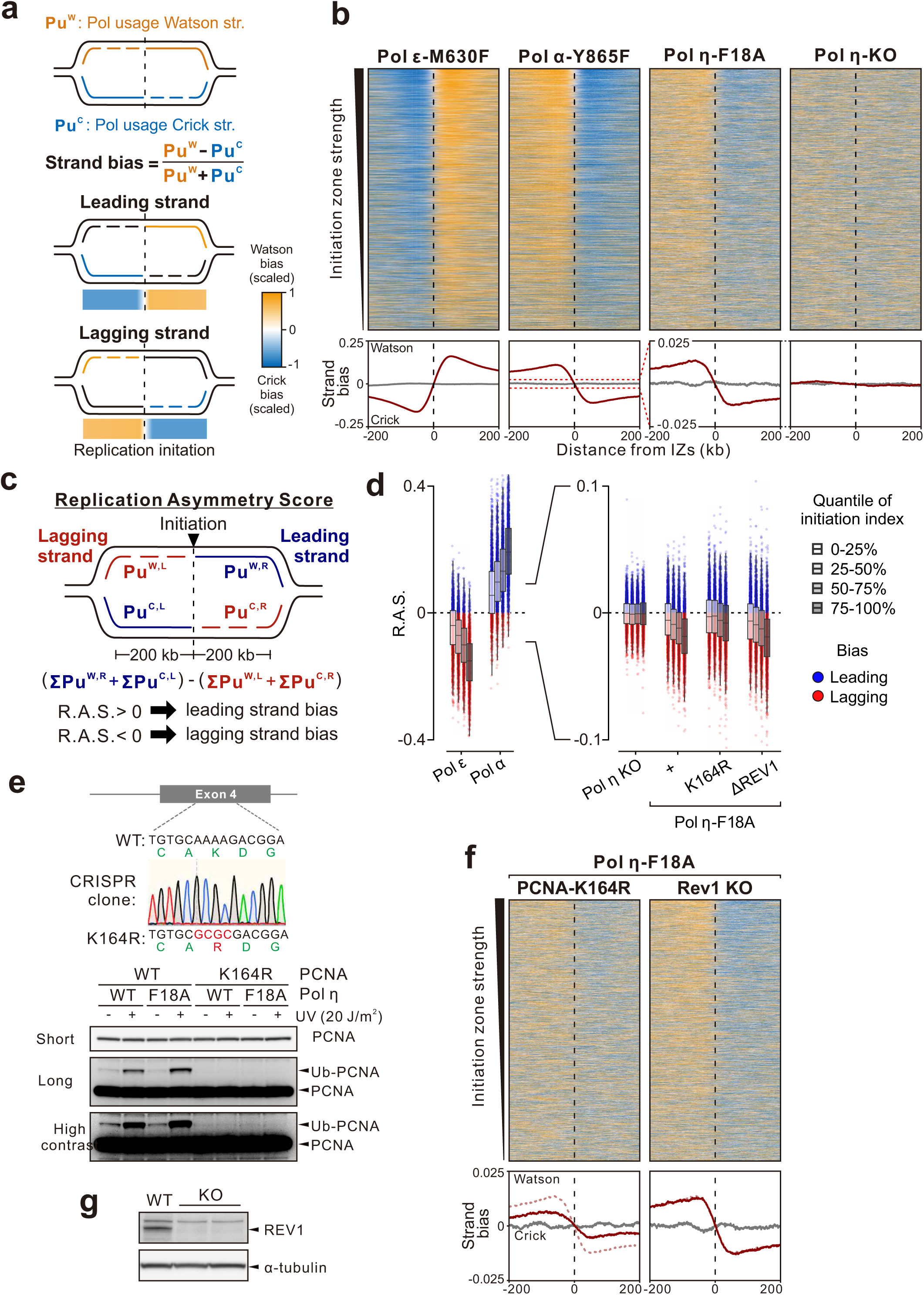
The Pol η usage profile displays a lagging strand bias. **a** Definition of strand bias score which quantifies the preferential usage of a polymerase on the Watson or Crick strand at each genomic position (see Materials and Methods for further details). Positive values (yellow) indicate Watson bias; negative values (blue) indicate Crick bias. **b** Top: Heatmap representation of strand bias scores for Pol ε, Pol α and Pol η usage ±200 kb around initiation zones (n = 7,139). The scaled strand bias was ranked by efficiency of replication initiation (top to bottom). Bottom: mean of strand bias around all initiation zones. **c** Definition of Replication Asymmetry Score (R.A.S.) which quantifies the replicative strand bias around each initiation zone (see Materials and Methods for further details). **d** R.A.S. of Pol η signal at individual initiation zones, categorised by replication initiation efficiency. **e** Top: sequencing confirming biallelic PCNA-K164R cells generated with CRISPR-Cas9. Bottom: ubiquitylated PCNA in parental cells and K164R mutant clones 8 hours after UV (20 J/m^2^) exposure. **f** Strand bias of Pol η usage around initiation zones in PCNA-K164R and Rev1-KO cells, plotted as in **b**. The dotted line in the bottom panel indicates the mean strand bias of Pol η usage without these regulatory mutations. **g** Western blot confirming REV1 knockout in WT Pol η (left) or Pol η-F18A cells (right).

To quantitatively relate Pol η activity to efficiency of replication initiation, we calculated a replication asymmetry score (R.A.S.; **Fig. 2c**) at each IZ. In this metric, positive values indicate a leading strand bias and negative values indicate a lagging strand bias. Determining R.A.S. at IZs categorised by efficiency of initiation (initiation index^19^) revealed increasingly negative R.A.S. values for Pol η at more efficient IZs, from which replication forks initiate more frequently in the cell population (**Fig. 2d**) This trend scaled in parallel with the asymmetry of Pol ε and Pol α usage, which served as the source data to generate the initiation index. The increased bias of Pol η, together with that of the replicative polymerases, clearly demonstrates the coupling of Pol η activity with replication forks.

Pol η is recruited to chromatin via PCNA, and ubiquitylation of PCNA at K164 further promotes Pol η function through an interaction with its ubiquitin-binding domain. To examine the contribution of this pathway to Pol η usage, we introduced a biallelic K164R mutation into the endogenous PCNA gene, preventing ubiquitylation at this residue under both unperturbed and UV-treated conditions (**Fig. 2e**). In these cells, Pol η Pu-seq profiles retained the lagging strand bias around IZs, but with reduced magnitude (**Fig. 2f, Supplementary** Fig. 6c), manifesting as a decrease in R.A.S. across IZs of all strengths (**Fig. 2d, Supplementary** Fig. 6d). Recruitment of Pol η can also be coordinated by the molecular scaffold REV1, which orchestrates polymerase switching during TLS^30^. We therefore examined the effect of REV1 on Pol η usage, by generating REV1-KO cells using CRISPR-Cas9 (**Fig. 2g**), but observed a similar strand bias of Pol η usage to REV1^+^ (**Fig. 2d,f, Supplementary** Fig. 6c). Together, these results indicate that the lagging-strand bias of Pol η is not entirely dependent on either canonical DNA damage tolerance regulatory pathway, though it is partially enhanced by PCNA ubiquitylation.

### Distinct Pol η usage across replication timing

In human cells, DNA replication is temporally coordinated, with early- and late-replicating regions distributed throughout the genome. Pu-seq profiles revealed regions with elevated Pol η signal, predominantly localised in late-replicating domains, which overlap with constitutive heterochromatin marked by H3K9me3 (**Fig. 3a, Supplementary** Fig. 7a). To assess this relationship genome-wide, we stratified the Pu-seq data into quantiles of Pol η signal intensity and examined their distribution relative to replication timing (RT). This analysis demonstrated a clear enrichment of high-intensity bins in late-replicating regions (**Fig. 3b, Supplementary** Fig. 7b), which was not observed in the Pol η-KO control. We performed the same analysis in PCNA-K164R cells, finding the late RT bias totally abolished, implicating PCNA ubiquitylation in this bias (**Fig. 3a,b, Supplementary** Fig. 7a,b). By contrast, Rev1 KO did not cause a substantial change in the distribution across RT (**Fig. 3b, Supplementary** Fig. 7a,b). These results implicate a critical role for PCNA ubiquitylation in mediating Pol η function during late replication.

**Figure 3.**
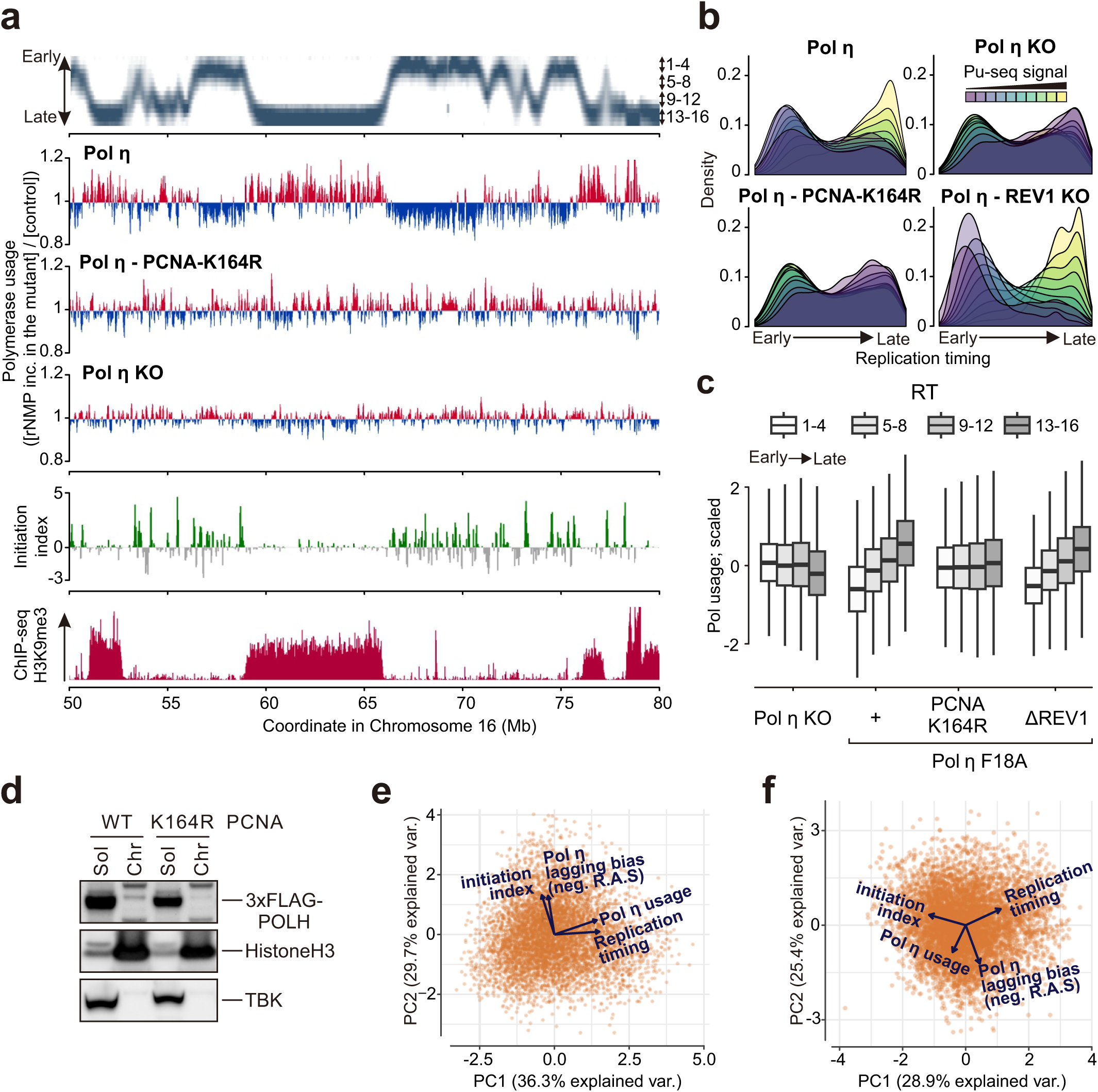
Usage of Pol η is bias towards late replicating regions. **a** Pol η usage across a representative region of Chromosome 16. Smoothed (*m* = 30), non-strand-specific Pu-seq profiles show enriched regions in red and depleted in blue. Replication timing (Zhao et al., 2020), initiation index (Koyanagi et al., 2022) and H3K9me3 ChIP-seq data are shown. Data visualised in IGV. **b** Density plots showing replication timing distribution of genomic bins stratified by Pol η Pu-seq signal intensity. **c** Boxplots show the scaled distribution of Pol η usage across replication timing (bins 1-4 = early; bins 13-16 = late). **d** Western blot of soluble and chromatin-bound fractions from cells with or without the PCNA-K164R mutation. **e** Biplots from PCA of Pol η Pu-seq signal properties at initiation zones genome-wide. **f** As **e**, but for Pol η Pu-seq in the PCNA-K164R background.

To test whether the late RT bias reflects changes in Pol η abundance, we monitored protein levels using biallelic FLAG-tagged Pol η cell lines. Following release from nocodazole-induced mitotic arrest, Pol η protein levels gradually increased as cells progressed through S phase (**Supplementary** Fig. 8a), consistent with a previous study^31^ and in line with the rising Pol η signal across replication timing (**Fig. 3c**). We next examined whether PCNA ubiquitylation influences chromatin association, by extracting the chromatin fraction. With WT PCNA, a small fraction of 3x-FLAG tagged Pol η is chromatin bound (**Figure 3d, Supplementary** Fig. 8b). Even in unperturbed conditions, PCNA ubiquitylation contributes to the enrichment of Pol η on chromatin, with significantly reduced chromatin-associated Pol η in PCNA-K164R cells. This effect was observed in both early and late stages of S phase (**Supplementary** Fig. 8c). We did not, however, detect an increase in the fraction of Pol η chromatin-bound between the stages of S phase. This may suggest that different PCNA ubiquitylation-dependent mechanisms, such as the recruitment of additional factors or alterations in the status of loaded PCNA, stimulate Pol η synthesis in these domains.

### The bias of Pol η toward the lagging strand and late-replicating regions represent two independent features

Our analyses indicate that Pol η usage is influenced by two distinct factors: (i) the structural asymmetry of DNA replication (leading vs. lagging strands) and (ii) megabase-scale replication timing domains. To test whether these two phenomena are functionally independent, we performed principal component analysis (PCA) using the R.A.S., initiation efficiency (initiation index), replication timing and total Pol η usage at all IZs. In the PCA biplot, initiation efficiency and R.A.S. showed similar loadings, with aligned vectors indicating a common underlying factor (**Fig. 3e**). By contrast, RT and total Pol η usage loaded together but orthogonally to initiation efficiency and R.A.S., suggesting that variation in Pol η usage across RT is functionally distinct from its strand bias at replication forks. This separation may reflect temporal changes in fork composition and polymerase engagement as replication progresses through S phase (see discussion). Repeating the PCA in PCNA-K164R cells, which lack the late-replication bias, revealed that total Pol η usage was no longer aligned with RT and R.A.S. was separated from initiation index vector (**Fig. 3f**). This weakened correlation between R.A.S. and initiation index reflects the partial contribution of PCNA ubiquitylation to lagging strand usage of Pol η. This analysis reinforces the conclusion that PCNA ubiquitylation influences the temporal and strand-specific usage of Pol η during unperturbed replication.

### Use of Pol η following UV damage maintains bias toward lagging strand

Since the most established role of Pol η is tolerating UV-induced DNA damage via TLS over CPD lesions, we next applied Pu-seq after UV exposure. To capture rNMP incorporation during lesion bypass, 5-ph-IAA was added to cells 4 hours before irradiation (5 J/m² UV), providing maximal degradation of RNaseH2A at the time of exposure (**Supplementary** Fig. 9a). Following this treatment, sufficient cell cycle progression was maintained to enable Pu-seq analyses, albeit with slower progression of S phase (**Supplementary** Fig. 9b). To mitigate the toxic effects of UV, cells were harvested after 24 hours.

CPDs form at dipyrimidine sites (TT, TC, CT, and CC), with TT lesions being the most prevalent^32^. Pu-seq detects the location of ribonucleotides at a 1bp-resolution, meaning the templating sequence context can be determined. To detect UV-dependent usage of Pol η, we compared the trinucleotide template context (the templating base and neighbouring 5′ and 3′ nucleotides) around ribonucleotides. As expected, UV exposure increased ribonucleotide incorporation upon TT template sequences (**Fig. 4a, Supplementary** Fig. 9c). The four most common template sequences for ribonucleotides after UV exposure were 3′-T<u>T</u>N-5′, corresponding to incorporation opposite the second thymidine of TT dinucleotides. Notably, the underrepresentation of 3′-N<u>T</u>T-5′ is unlikely to reflect a biological preference of Pol η for the second base over the first. Rather, Pu-seq is more likely to detect rNMP incorporation opposite the second base, since, when multiple rNMPs are incorporated, Pu-seq detects the position of the 3′-most (e.g. when two rNMP are sequentially incorporated, Pu-seq detects the second). Importantly, detecting elevated Pol η usage precisely at predicted damage sites directly links our Pu-seq signal to polymerase activity in cells.

**Figure 4.**
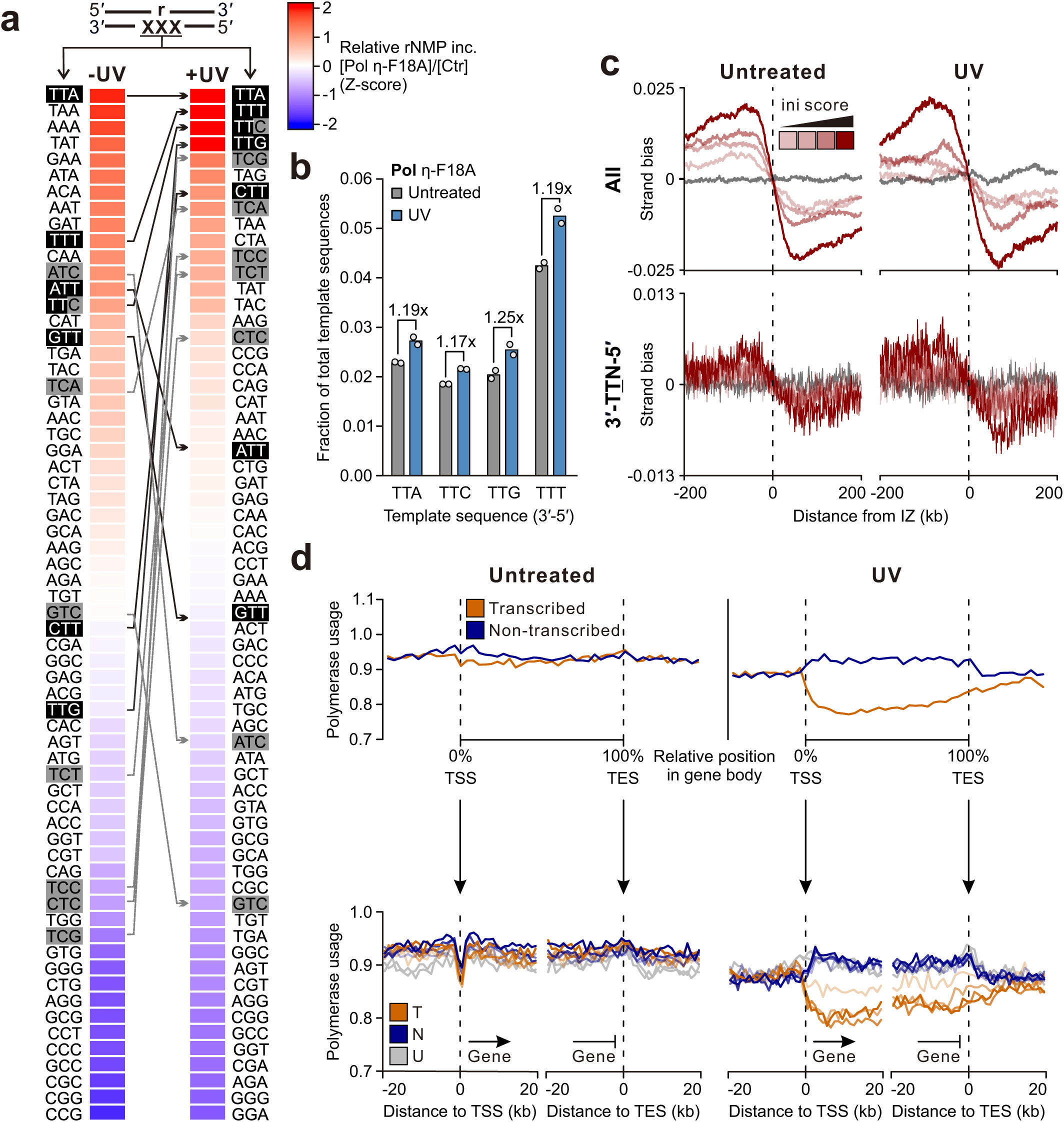
Altered distribution of Pol η usage after UV damage. **a** Relative frequency of ribonucleotides detected at each trinucleotide template sequence in Pol η-F18A cells vs control cells, with and without 5 J/m^2^ UV. Ratios were calculated for each trinucleotide, z normalised and sorted by Z-score. Template sequences are displayed 3′ to 5′, with CPD-forming TT’s in black and TC’s in grey. **b** Fraction of total detected ribonucleotides from Pol η-F18A cells located at CPD-forming sequences, with and without 5 J/m^2^ UV. **c** Aggregate strand bias of Pol η usage aligned at IZ, sorted by strength. Smoothed (*m* = 30) profiles are shown for all detected ribonucleotides and those deriving from the top CPD-forming sequences (3′-T<u>T</u>N-5′), with and without UV. **d** (Top) Usage of Pol n on the transcribed or non-transcribed strand along scaled gene bodies for the longest 50% of genes among the top 50% of expressed genes (RNA-seq FKPM; Koyanagi et al, 2022), with and without UV. (Bottom) Aggregate usage profiles aligned at the start and end sites of all genes, quartiled by expression level, with opacity reflecting expression. T = Transcribed strand, N = Non-transcribed strand, U = Inactive gene.

Following UV exposure, the detection of ribonucleotides at the top four enriched TT sequences increased by ∼20%, to a total of 12.7% of Pu-seq reads and indicating only a minor contribution relative to Pol η usage during unperturbed replication (**Fig. 4b**). Consistent with this, the strand bias of the Pol η signal around efficient initiation zones remained comparable to untreated cells (**Fig. 4c, top, Supplementary** Fig. 9d). No noticeable change in strand bias was observed at the enriched TT dinucleotides themselves either (**Fig. 4b, bottom**). Thus, under these conditions, TLS by Pol η did not override its intrinsic replication-associated bias, which remained the dominant signal.

Although strand bias remained similar, UV exposure caused a reduction in the overall Pol η Pu-seq signal around IZs (**Supplementary** Fig. 9e**, top**). We previously identified that IZs are commonly flanked by genes^19^. Interestingly, transcriptional activity around IZs displays an almost exact inverse relationship with the UV-dependent reduction in Pu-seq signal (**Supplementary** Fig. 9e**, bottom**). We therefore hypothesised that transcription-coupled nucleotide excision repair (TC-NER), which selectively removes DNA lesions from the template strand of RNA polymerase, may underlie the reduction. Consistent with this, UV exposure induced a pronounced bias of Pol η usage towards the non-transcribed strand within gene bodies, which was not observed in unperturbed conditions (**Fig. 4d**). This bias began precisely at transcription start sites (TSS) and extended ∼10 kb downstream of transcription end sites (TES), likely reflecting transcriptional readthrough, where RNA polymerase continues transcribing for some distance beyond the TES^33^. The absence of a transcriptional bias under normal conditions indicates that Pol η activity is not driven by bulky lesion bypass, but instead reflects its broader role in DNA replication. Together, our data show that UV induces a detectable reduction in Pol η-mediated TLS on the transcribed strand, but no detectable shift in fork-oriented bias under both unirradiated and irradiated conditions (see Discussion).

### Usage of Pol η in a non-cancer cell line

Human Pu-seq was originally developed in the colorectal cancer cell line HCT116. However, cancer cells often exhibit replication stress due to disruptions in DNA replication pathways. To determine whether our findings reflect general features of replication, rather than cancer-specific adaptations, we established an analogous Pu-seq system in immortalised, non-transformed RPE1 cells. Pu-seq profiles for Pol ε, Pol α and Pol η were generated, which broadly correlated with their respective datasets from HCT116 (**Fig. 5a,b**). In RPE1 cells, Pol α and Pol η usage was particularly highly correlated, and the lagging strand bias of Pol η was even more pronounced than in HCT116 (**Fig. 5c**). Pol η also retained a bias towards late-replicating regions (**Fig. 5d**). These results suggest these characteristics of Pol η usage are conserved in non-transformed human cells and likely represent an intrinsic feature of human DNA replication.

**Figure 5.**
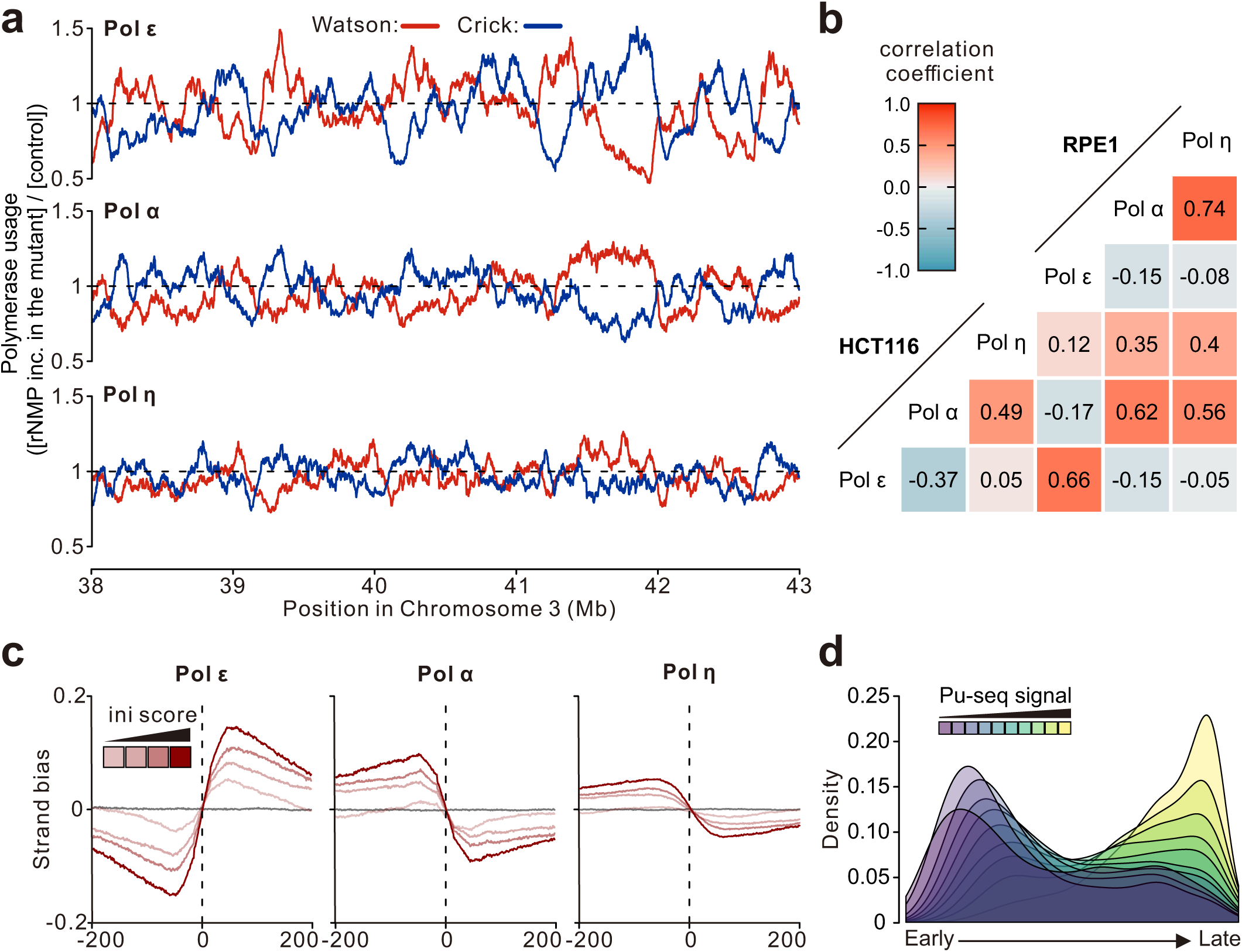
Polymerase usage patterns in non-transformed RPE-1 cells. **a** Smoothed (*m* = 30), strand-specific polymerase usage of Pol ε, Pol α and Pol η across a representative region of Chromosome 3. Red: Watson strand. Blue: Crick strand. **b** Heatmap of Pearson correlations between genome-wide strand-specific profiles of Pol ε, Pol α and Pol η variants in HCT116 and RPE1 cells. Red: positive. Blue: negative. **c** Mean strand bias of Pol ε, Pol α and Pol η around IZ (n = 4,874), divided into quartiles by initiation strength. **d** Density plots showing replication timing (Zhao et al., 2020) distribution of genomic bins stratified by Pol η Pu-seq signal intensity.

### Errors by Pol η contribute to single nucleotide mutations in cancers

Error-prone polymerases are inherently low-fidelity, and Pol η is particularly prone to introducing A to G substitutions by misincorporating dGTP opposite template T^34^. Given its preferential activity on specific strands and within particular genomic contexts, we asked whether Pol η usage patterns might be reflected in the mutational landscapes of human tumours. Analysis of The Cancer Genome Atlas (TCGA) datasets revealed that, in several cancer types, such as gastric adenocarcinoma, A to G mutations exhibit strand asymmetry around replication IZs consistent with enrichment on the lagging strand (**Fig. 6a**) and a marked enrichment within late-replicating regions relative to the overall mutation distribution (**Fig. 6b**).

**Figure 6.**
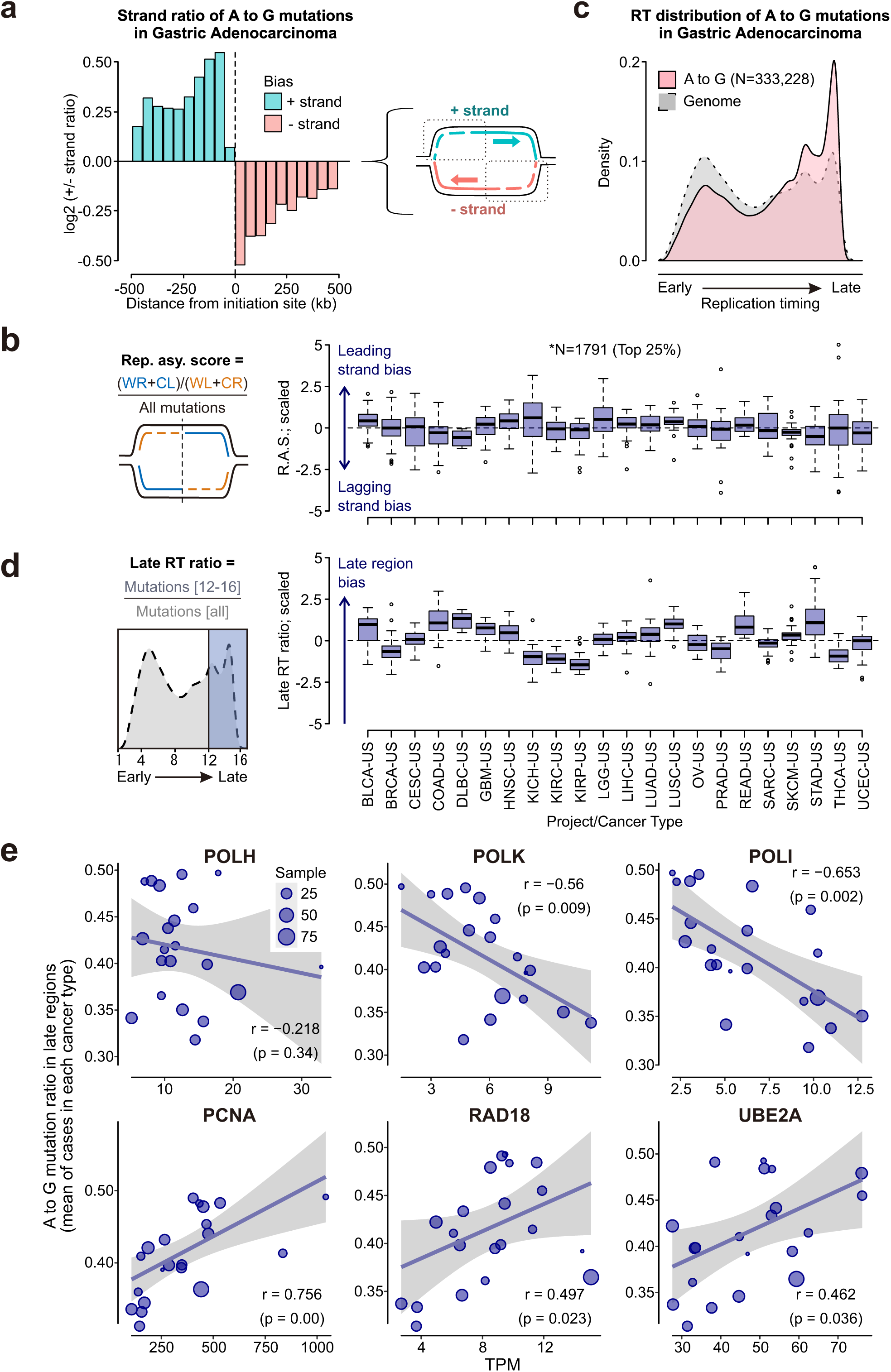
Distribution of characteristic Pol η mutations in human cancers from the TCGA database. **a** Strand asymmetry of A to G mutations in gastric adenocarcinoma, aligned at the strongest 25% of IZ, plotted as log2(Watson/Crick) mutation ratio. **b** Average R.A.S. of A to G mutations in individual patients, scaled and grouped by cancer type. **c** Distribution of A to G mutations relative replication timing (Zhao et al., 2020). **d** Ratio of A to G mutations in late-replicating regions (bins 12-16) relative to all regions in individual patients, scaled and grouped by cancer type. **e** Correlation of gene expression (TPM) and the ratio of A to G mutations occurring in late-replicating domains. Points represent mean values for each cancer type.

To investigate mutational distribution more thoroughly, we quantified strand bias at the individual-patient level within the groups. Calculating R.A.S. for A to G mutations revealed an overall lagging strand bias in several cohorts (**Fig. 6c**), though the scaled data indicated only minor inter-type variation. By contrast, the ratio of A to G mutations in late-replicating regions varied substantially across the cohorts, revealing a pronounced late-replication bias in several cancer types (**Fig. 6d**).

Given this variation in late-replication bias, we next asked whether these patterns were associated with expression of genes related to Pol η function. To investigate this, we correlated mRNA expression levels with the mutational distributions. *POLH* expression itself did not correlate with the A to G mutation enrichment in late-replicating regions across cancer types, when taking the mean of each cohort (**Fig. 6e**), consistent with previous work showing that overexpression of Pol η alone does not increase spontaneous mutation rates^35^. In contrast, the expression of *PCNA* and both *RAD18* and *UBE2A* (the E3 and E2 ligases responsible for PCNA-K164 ubiquitylation) positively correlated with enrichment of A to G mutations in late replicating regions, consistent with PCNA ubiquitylation-dependent Pol η synthesis at these sites. Importantly, *RAD18* and *UBE2A* expression also correlated significantly with all mutations from A/T bases, but not from C/G bases (**Supplementary** Fig. 10), reflecting the known mutational preferences of Pol η synthesis observed in vitro^34^. Interestingly, expression of Y-family polymerases Pol κ and Pol ι, which also bind to ubiquitylated PCNA, negatively correlated with A to G mutation enrichment in late-replicating regions. As these polymerases can be regulated in a transcription-dependent manner^36, 37^ and Pol κ has been implicated in early S phase replication^38^, competition among error-prone polymerases may influence the timing of mutation occurrence and its genomic distribution. Altogether, these analyses suggest that PCNA ubiquitylation plays a role in late-replication-associated A to G mutagenesis, consistent with Pol η activity, and that the balance between Y-family polymerases may contribute to mutational patterns across S phase.

## Discussion

Here we establish a method to directly monitor Pol η synthesis in human cells and identify two features of genome replication that correlate with the Pol η function under unperturbed conditions: (i) lagging strand replication and (ii) replication timing. While the former has been observed for the budding yeast Pol η^20^ (Rad30) and the *E. coli* homolog (Pol V) under SOS response^39^, this work provides the first evidence that the lagging strand bias is conserved in mammalian cells. These findings suggest that association of Pol η with the lagging strand is either an evolutionary conserved property or an inherent consequence of discontinuous lagging strand synthesis. In contrast, the bias towards late-replicating regions appears to be specific to large and complex genomes, since such domains are absent in compact genomes, underscoring the importance of studying replication in mammalian systems to capture the full complexity of eukaryotic replication dynamics.

### Stochastic Pol η usage at replication forks

The eukaryotic replisome exhibits substantial heterogeneity due to the stochastic nature of multiprotein assemblies providing opportunities for concentration-dependent polymerase exchange^40^. To evaluate potential roles for Pol η in lagging strand synthesis, we consider how intrinsic mechanisms of leading and lagging strand replication influence polymerase usage. As a core component of the replisome, Pol ε is directly coupled to the CMG helicase. In contrast, Pol δ maintains only a weak association with DNA and can readily dissociate from PCNA^41, 42^. Because PCNA dissociation from DNA is relatively slow^41^, Pol δ release can leave a PCNA-bound primer-template end, a potential substrate for error-prone polymerases (**Fig. 7**). Pol η, which travels with the replication fork^12^, interacts with PCNA through three redundant PIP-box motifs^43^ and may compete for access to PCNA-bound sites, which could arise on a subset of Okazaki fragment termini. Importantly, PCNA-bound Pol η was shown to promote PCNA ubiquitylation, which in turn enhances local accumulation of Pol η^43^. Together with studies showing that Okazaki fragments can trigger PCNA ubiquitylation to promote their synthesis and maturation^44, 45, 46^, our finding that PCNA ubiquitylation enhances Pol η lagging-strand bias suggests a local mechanism that increases Pol η recruitment. Moreover, since Pol η can extend RNA–DNA junctions in vitro, it might initiate synthesis from RNA primers in vivo following inefficient primase-Pol α handoff^47^ (**Fig. 7**). Thus, elevated polymerase turnover on the lagging strand may increase the opportunity for error-prone polymerases, such as Pol η, to stochastically engage and extend exposed PCNA-bound DNA ends.

**Figure 7.**
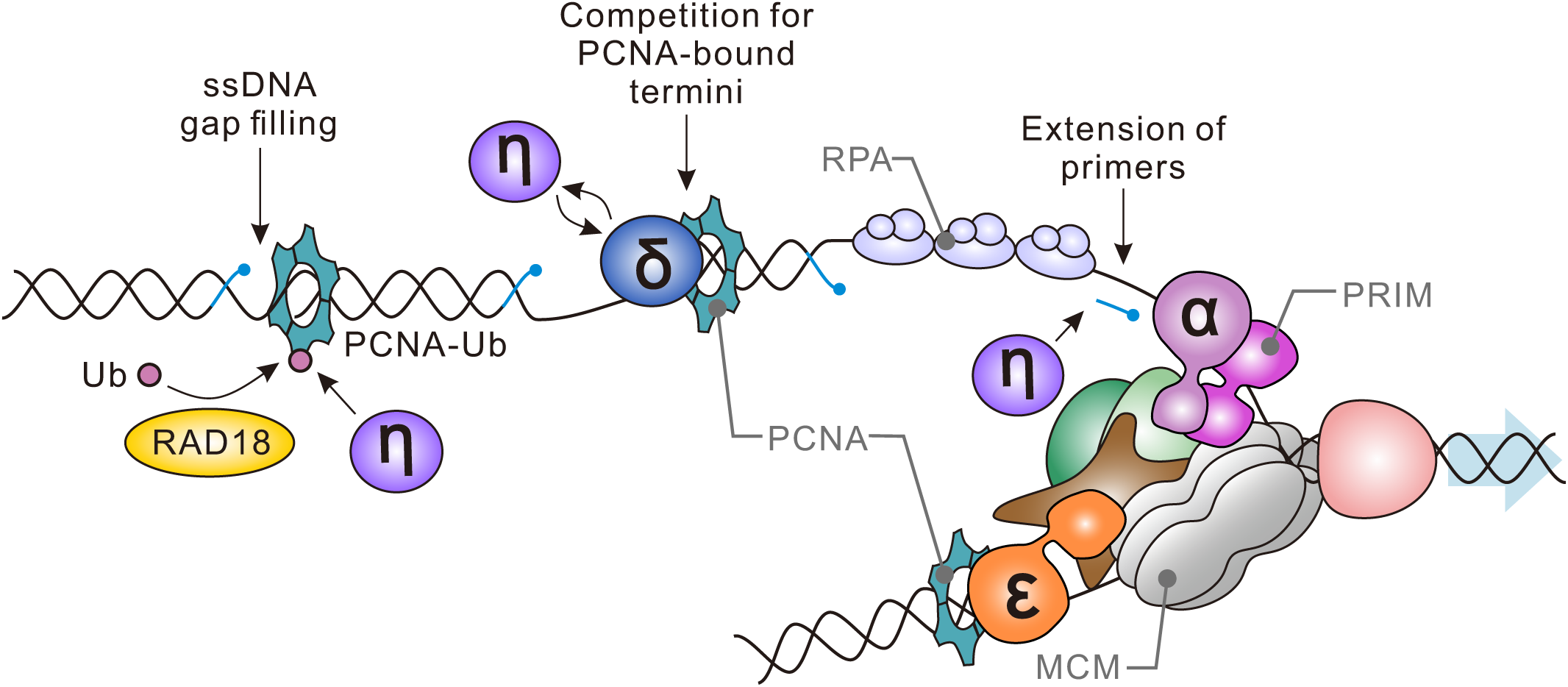
Mechanism of Pol η recruitment during lagging strand synthesis. At the replication fork, the MCM helicase complex (grey) unwinds DNA. Pol ε (orange) synthesises the leading strand continuously while tethered to the MCM. On the lagging strand, Pol α-primase (pink) synthesises short RNA-DNA primers (light blue), which Pol δ (blue) usually extends. Pol η (purple) may also extend from RNA or DNA primers. Pol δ and Pol η both associate with PCNA (turquoise), suggesting Pol η can compete with Pol δ for binding and access to the lagging strand termini. Formation of ssDNA gaps between Okazaki fragments triggers PCNA ubiquitylation (dark blue), enhancing Pol η recruitment for gap filling.

However, it remains unclear whether Pol η actively serves as a designated “backup” for Pol δ or whether its usage simply reflects stochastic polymerase dynamics. Notably the Pol η-KO HCT116 cells we generated showed no overt defects, which may indicate that other error-prone polymerases (e.g. Pol κ, Pol ι or Pol ζ) can redundantly support Okazaki fragment synthesis. The requirement for Pol η and Pol κ increases upon cell cycle dysregulation and replication stress, such as that induced by oncogene activation, which can elevate fork stalling and PCNA ubiquitylation^48, 49^. Under such conditions, the backup functions of error-prone polymerases become critical for sustaining efficient progression of replication. As a future direction, we are extending Pu-seq to various error-prone polymerases to define how their activities are coordinated under replication stress in human cells.

### Pol η usage under unperturbed conditions is unlikely to reflect lesion bypass

Stochastic dynamics at replication forks provide a simple explanation for Pol η engagement during unperturbed replication, but an alternative is that the observed lagging-strand bias reflects asymmetric tolerance of endogenous lesions. Analyses of large-scale mutagenesis in human genomes revealed elevated instances of error-prone synthesis on the lagging strand and similar mutagenic spectrums on the lagging and non-transcribed, suggesting a shared error-prone bypass pathway in both contexts^50^. In our study, Pol η activity exhibited a clear transcriptional strand bias only after UV irradiation, with no bias observed under unperturbed conditions (**Fig. 5d**). This pattern implies that, under normal growth conditions, DNA lesions that are recognised by TC-NER are not present and Pol η activity largely reflects synthesis on undamaged templates. We therefore favour the interpretation that the lagging-strand bias of Pol η usage is largely independent of DNA damage tolerance and does not imply biased usage during DNA damage bypass. Consistent with this interpretation, damage–induced mutagenesis during mouse liver tumorigenesis produces TLS-dependent clustered mutations with no discernible leading and lagging strand asymmetry^51^.

Despite observing transcriptional strand asymmetry after UV exposure, we detected no substantial change in replicative strand bias of Pol η synthesis relative to untreated cells. We interpret this as evidence that, under mild UV exposure (5 J/m², a dose that permits continued cell cycle progression), TLS over UV-induced lesions constitutes a minor contribution versus the predominant replication-associated Pol η signal. Taken together, our data support a model in which Pol η usage is largely governed by the stochastic architecture of replication forks.

### PCNA ubiquitination-dependent polymerase usage contributes to mutagenesis during late replication

Pol η usage was biased towards late-replicating regions of the genome enriched with the repressive histone modification H3K9me3, characteristic of constitutive heterochromatin. These regions lack defined replication initiation zones and do not exhibit defined replication fork directionality^19, 29^. In our previous Pu-seq analyses with replicative polymerases Pol ε and Pol α, these regions showed no reciprocal strand asymmetry^19^. While this could represent random initiation of DNA replication, it could also reflect distinct polymerase assignment relative to other genomic domains.

In addition to accessibility, late-replicating heterochromatin differs from early-replicating euchromatin in several respects. For instance, replication forks progress more rapidly through these domains^52^. Elevated MCM unwinding rates at increased fork speeds may increase the spatial separation between lagging strand priming and synthesis, raising the likelihood of synthesis becoming uncoupled from the replisome. Since PCNA ubiquitylation has been implicated in post-replicative gap filling^53, 54^, such uncoupling could leave ssDNA gaps that recruit Pol η in a PCNA ubiquitylation– dependent manner (**Fig. 7**). Supporting this model, recent single-molecule imaging in unperturbed cells revealed chromatin-associated Pol η spatially distinct from sites of active DNA synthesis, with increased foci observed in cells with high DNA content (i.e., late S or G2 phase)^55^. Moreover, error-prone Pol ζ (REV3L) has been implicated in facilitating replication through heterochromatin^56^, suggesting cooperative roles for these polymerases in mitigating under-replication (ssDNA gaps, etc.) in compact, late-replicating chromatin domains. Although the precise molecular mechanisms remain to be defined, accumulating evidence supports the notion that the replication dynamics varies over the course of S phase in human cells.

Consistent with the PCNA ubiquitylation–dependent Pol η usage in late-replicating regions, we found a positive correlation between the mutation burden in these domains in cancer genomes and the expression of *RAD18* and *UBE2A*, key components of the PCNA ubiquitination pathway. Late-replicating regions are associated with high mutation loads in various biological systems (e.g., comparisons of human and chimpanzee genomes^57^, analyses of lymphoblastoid cell lines^58^), yet the underlying mechanisms remain incompletely understood. Our analysis of cancer mutations revealed that the most common substitutions made by Pol η were often enriched in late-replicating regions, where the frequency correlates with *RAD18* expression (**Supplementary** Fig. 10). We therefore propose RAD18-dependent PCNA ubiquitylation as one of the mechanisms that contributes to the elevated mutation load in late-replicating domains. Importantly, variation of *RAD18* expression also correlates with tumour aggressiveness and poor clinical outcomes (reviewed in ^18^). Given the critical role of PCNA ubiquitylation in post-replication repair, the resulting mutations may represent an unavoidable trade-off of this “last resort” mechanism for ensure timely completion of genome replication. Future studies will define and characterise the broader mutational spectrum arising from error-prone Pol η synthesis.

### Comparison with other methodologies

Detecting polymerase usage with Pu-seq offers a distinct advantage, as it captures nucleotide incorporation directly in vivo, generating high-resolution, strand-specific maps of genome-wide activity. Here, we demonstrate that Pu-seq is sufficiently sensitive to detect the activity of even low-abundance polymerases such as Pol η, a sensitivity that is difficult to achieve with conventional methods. For example, our attempt to profile Pol η enrichment in late-replicating regions by ChIP-seq of endogenously tagged 3×FLAG-POLH failed to detect enrichment over the untagged control. This likely reflects a combination of Pol η’s low abundance, its transient or weak DNA association and its lack of strong site-specific localisation (unpublished data). These limitations highlight the advantage of directly measuring DNA synthesis events for detecting low-abundance polymerase activity in vivo. A deeper understanding of the roles of these polymerases will elucidate the mechanisms safeguarding genome stability and may help explain how the mutational landscape is shaped during human genome replication.

## Materials and Methods

### Protein purification

Hexahistidine tagged Pol η at N-terminus (his-Pol η) and its F18A mutant were purified as described previously^59^.

### Primer extension assays

Nucleotide sequences of oligonucleotide templates and the primer are provided in Supplementary Table 1. The CPD-containing template^60^ was purchased from Operon Biotech. The primer was labelled using polynucleotide kinase (New England Biolabs) and [γ-^32^P] ATP (Revvity Health Sciences, Inc. BLU502Z) and subsequently annealed to the templates. The standard reaction mixture (10 μL) contained 20 mM HEPES– NaOH (pH 7.5), 50 mM NaCl, 0.2 mg/ml bovine serum albumin (BSA), 5 mM DTT, 5 mM MgCl_2_, 100 mM of each dNTP, 1 pmol primer/template and the indicated amounts of his-Pol η. Reactions were incubated at 30°C for 5 mins and terminated by adding 10 μL of stop solution (30 mM EDTA, 94% formamide, 0.05% bromophenol blue, 0.05% xylene cyanol). Reaction products were resolved on 8% polyacrylamide gels containing 7 M urea and visualized and analysed using a Typhoon FLA 7000 (GE Healthcare). Kinetic assays were performed at 30 °C for 5 mins in 10 μL standard reaction mixture in the presence of one dNTP or rNTP at the indicated concentrations. Reaction products were resolved on 20% polyacrylamide gels containing 7 M urea. *K_m_* and *k_cat_* were determined from plots of initial velocity versus NTP concentration, obtained from three or four independent experiments using a hyperbolic curve-fitting program (OriginLab Corporation).

### Cell culture

Cell lines derived from HCT116 cells (ATCC #CCL-247) or hTERT RPE-1 (ATCC #CRL-4000) were cultured in McCoy’s 5A (modified) Medium (Gibco) or DMEM/F-12 (Gibco), respectively, supplemented with 10% fetal bovine serum (FBS, Gibco), 2mM L-glutamine, 100 U/ml penicillin and 100 μg/mL streptomycin. Cells were maintained at 37 °C in a humidified atmosphere containing 5% CO_2_. 5-ph-IAA (BioAcademia) was prepared as a 250 μM stock solution in dimethyl sulfoxide (DMSO) and stored at - 20 °C. dTAG^V^-1 (R&D Systems) was prepared as a 500 μM stock in DMSO and stored at -20 °C. Drugs were added directly to the culture medium to achieve final working concentrations of 0.4 μΜ (5-ph-IAA) and 0.5 μM (dTAG^V^-1).

### Generation of endogenous gene mutations

CRISPR-associated (Cas) proteins (Alt-R^TM^ A.s. Cas12a (Cpf1) Ultra or Alt-R^TM^ S.p. HiFi Cas9 Nuclease V3) were purchased from IDT (Integrated DNA Technologies, USA). For the POLH-F18A mutation, ribonucleoprotein (RNP) complexes were assembled by combining equal volumes of 63 μΜ Cas12a protein and 75 μM Alt-R^TM^ A.s. Cas12a crRNA (IDT; Custom production; Supplementary Table 1). For the PCNA-K164R mutation and tagging of POLH with 3xFLAG, guide RNA (gRNA) molecules were assembled by mixing 50 μM Alt-R^TM^ CRISPR-Cas9 crRNA (IDT; Custom production; Supplementary Table 1) with 50 μM Alt-R™ CRISPR-Cas9 tracrRNA (IDT) in Nuclease-Free Duplex Buffer (IDT), heating at 96 °C for 5 mins, then cooling to RT. RNP complexes were assembled by combining 125 pmol Cas9 enzyme with 150 pmol gRNA. For electroporation, 1 x 10^5^ cells were mixed with 1.5 μL RNP and either 1.5 μL single-stranded Alt-R™ HDR Donor Oligo (IDT; Custom production; Supplementary Table 1) or 2 μL 500 nM double-stranded Alt-R™ HDR Donor Block (IDT; Custom production; Supplementary Table 1) to 15 μL in R buffer. Electroporation was performed using the Neon Transfection System (Invitrogen) under the following conditions: HCT116: one pulse of 1,530 V for 20 ms; RPE1: two pulses 1,350 V for 20 ms. Cells were plated in media containing 0.33 μM Alt-R™ HDR Enhancer V2 (IDT) for 24 hours, then media was replaced and cells were incubated for a further 48 hours at 37 °C to allow recovery. Subsequently, individual clones were isolated, expanded and screened for the presence of the genetic modifications.

### Generation of gene knockout mutations

A frameshift mutation was introduced into genes using the CRISPR-Cas9 system. Specifically, Cas9 and a gRNA (Supplementary Table 1) targeting POLH (exon 3, amino acid 83/713) or REV1 (exon 5, amino acid 232/1250) were expressed from the pSpCas9 vector^61^. The plasmid was transfected into cells using FuGeneHD (Promega) following the manufacturer’s instructions. After 24 hrs, puromycin (Gibco) was added to a final concentration of 0.25 μg/mL to select for transfected cells. Upon complete selection, individual clones were isolated, expanded and screened for the presence of knock-out mutations. Gene knock-outs were verified by western blotting.

### Cell growth assay

Cells were seeded in a 6-well plate and allowed to adhere for 48 hrs. Following attachment, cells were treated with 0.4 μΜ 5-ph-IAA (BioAcademia, Japan, #30-003) where required. At the indicated timepoints, cells were harvested and counted using a Countless 3 Automated Cell Counter (Invitrogen). To assess relative cell growth, cell counts at each time point were normalized to the cell number at 0 hrs.

### Cell survival assay

Cells (2 x 10^3^ per well) were seeded in a 6-well plate and allowed to adhere for 48 hrs. Following attachment, cells were either exposed to the indicated dose of UV radiation using a crosslinker (UVP CX-2000) or continuously grown in 0.4 μM 5-ph-IAA. Colonies were allowed to form for approximately 10 (UV) or 7 (5-ph-IAA) days, with culture medium and drugs replaced as needed. To assess cell survival, colonies were stained with a 0.5% crystal violet solution (0.5% crystal violet in 20% methanol) for 20 mins. Excess stain was removed by washing with Milli-Q H_2_O and plates were dried overnight. For quantification, the crystal violet stain was dissolved in 33% acetic acid and absorbance at 590 nm measured using a plate reader (Thermo Scientific). Relative cell survival was calculated by comparing the treated conditions to the untreated control.

### Cell synchronisation with nocodazole

Cells (1 x 10^5^ per well) were seeded in a 6-well plate and allowed to adhere for 48 hrs. Synchronisation in G2-phase was achieved by treatment with 50 ng/mL nocodazole for 15 hrs. To release cells from arrest, cultures were washed twice with fresh medium. Samples were harvested at the indicated timepoints and processed for analysis by western bott and flow cytometry.

### Chromatin extraction

Cells were harvested and washed once with ice-cold PBS. The pellet was resuspended by gentle pipetting in 50 µL of 0.2% Triton X-100 in PBS supplemented with protease inhibitor and incubated on ice for 5 mins with occasional gentle mixing. Lysates were centrifuged at 17,800 g at 4 °C for 10 mins and the supernatant was collected as the soluble fraction. The pellet was resuspended in 200 µL of 0.2% Triton X-100 in PBS with protease inhibitor, underlayered with 200 µL of 30% (w/v) sucrose and centrifuged at 17,800 g at 4 °C for 10 min. The supernatant was carefully aspirated and the sucrose separation step repeated. The final pellet was resuspended in 50 µL of 0.2% Triton X-100 in PBS with protease inhibitor and treated with benzonase on ice for 15 mins to yield the chromatin fraction. Samples were mixed with 4× Bolt LDS Sample Buffer and 10× Bolt Reducing Agent and heated for 10 mins at 70 °C.

### Western blotting

Cells were lysed in RIPA buffer (Sigma-Aldrich) supplemented with protease inhibitors and sonicated (30 sec on, 30 sec off for 15 cycles) using a Bioriptor II (BMBio). Following sonication, lysates were centrifuged at 10,000 g for at 4 °C 20 mins. The supernatant was collected and mixed with 4x Bolt^TM^ LDS sample buffer (Invitrogen™) and 10x Bolt^TM^ Sample Reducing Agent (Invitrogen™). Samples were incubated at 70 °C for 10 mins and proteins separated by SDS-PAGE on Bolt^TM^ 8% or 4-12% Bis-Tris Plus gels (Invitrogen). Following electrophoresis, proteins were transferred to Immun-Blot PVDF membranes (BioRad) using a Power Blotter XL (Invitrogen). Membranes were blocked with 5% milk in phosphate buffered saline (PBS) containing 0.1% Tween-20 (PBST). Membranes were incubated with primary antibodies overnight at 4 °C (anti-RNaseH2A, anti-PolH, anti-FLAG, anti-PCNA or anti-REV1) or for 1 hr at room temperature (anti-tubulin) in 0.5% milk in PBST. Secondary antibody incubations were performed at room temperature for 1 hr in 0.5% milk in PBST. Detection was conducted using Western Lightning® Plus-ECL (Perkin Elmer) and images were acquired with a ChemiDoc^TM^ Touch MP Imaging system (Bio-Rad).

The following antibodies were used: anti-RNASEH2A (dilution 1:2,000, Bethyl Laboratories® A304149A), anti-PolH (dilution 1:4,000, Bethyl Laboratories® A301-231A), anti-PCNA (dilution 1:1,000, Sigma-Aldrich MAB424), anti-REV1 (dilution 1:2,000, Santa Cruz Biotechnology sc-393022), anti-FLAG (dilution 1:2,000, Sigma-Aldrich F1804), anti-tubulin (dilution 1:5,000, Sigma-Aldrich T5168), anti-Histone H3 (dilution 1:2,000, Abcam ab1791), anti-NAK1/TBK (dilution 1:2,000, Abcam ab40676), anti-mouse IgG HRP conjugate (dilution 1:10,000, Agilent P026002-2) and anti-rabbit IgG HRP conjugate (dilution 1:10,000, Agilent P044801-2).

### Flow cytometry

Cells (2 x 10^4^ per well) were seeded in a 6-well plate and allowed to adhere for 48 hrs. Following attachment, 0.4 μM 5-ph-IAA was added where required. After an additional 48 hrs, cells were harvested, fixed in 70% ethanol and stored at -20 °C for 1-7 days. Prior to analysis by flow cytometry, cells were washed with PBS and resuspended in PBS containing 1% BSA, 40 μg/ml propidium iodide (PI), and 50 μg/ml RNaseA. The suspension was incubated for 30 mins at 37 °C, then on ice for two minutes. Samples were analysed on a BD Accuri^TM^ C6 Plus Flow Cytometer (BD Biosciences) and approximately 1 x 10^5^ cells were analysed using FlowJo^TM^ software (BD Biosciences).

### ChIP-seq

Semi-confluent cells in 10 cm dishes were crosslinked in 1.42% formaldehyde (final concentration) in PBS for 15 mins at room temperature with gentle agitation. Crosslinking was quenched with 127 mM glycine for 5 minutes. Cells were washed with PBS, lysed in buffer (10 mM Tris-HCl pH 8, 10 mM NaCl, 0.5% IGEPAL® CA-630), and pelleted (2,000 g, 5 mins, 4 °C). Pellets were resuspended in SDS lysis buffer (50 mM Tris-HCl pH 8, 1% SDS, 10 mM EDTA), followed by addition of ice-cold RIPA buffer (50 mM HEPES, 1 mM EDTA, 140 mM NaCl, 1% Triton X-100, 0.1% Sodium deoxycholate) supplemented with protease inhibitors (Roche Complete and 1 mM AEBSF).

Chromatin was sheared using a Bioruptor (30 sec on, 30 sec off for 60 cycles), yielding ∼330 μL clarified lysate per sample after centrifugation (12,000 g, 10 min, 4 °C). A 30 μL aliquot was reserved as input. Immunoprecipitation was carried out using 2 μg of anti-Histone H3K9me3 antibody (Abcam ab8898) per 300 μL chromatin, incubated for 3 hrs at 4°C with rotation, followed by addition of 20 μL pre-washed Protein G Dynabeads and further incubation for 1 hr. Beads were sequentially washed with RIPA buffer, wash buffer 1 (50 mM HEPES, 1 mM EDTA, 500 mM NaCl, 1% Triton X-100, 0.1% Sodium deoxycholate), wash buffer 2 (10 mM Tris-HCl (pH8), 0.25 M LiCl, 0.5% IGEPAL® CA-630, 0.5% Sodium deoxycholate, 1 mM EDTA), and TE, each with rotation at 4°C. Elution was performed in elution buffer (50 mM Tris-HCl (pH8), 10 mM EDTA, 1% SDS) at 65°C for 15 mins, followed by overnight reverse crosslinking at 65°C.

Input and immunoprecipitated DNA samples were treated with RNase A and Proteinase K, extracted with phenol/chloroform, and precipitated with ethanol in the presence of sodium acetate and glycogen. DNA pellets were washed with 80% ethanol, air-dried and resuspended in 10 mM Tris-HCl pH 7.5. DNA concentrations were determined with PicoGreen and libraries were prepared using the NEBNext Ultra II DNA Library Prep Kit for Illumina according to the manufacturer’s protocol. Libraries were sequences as 150-bp paired-end (PE) reads on an Illumina Hiseq X platform (Macrogen, Tokyo, Japan). One biological replicate was analysed.

### Pu-seq assays

Standard Pu-seq assays were conducted as described previously^19^. Briefly, 1.25 x 10^4^ cells/ml were seeded in 15 cm petri dishes (Greiner). After 48 hrs, RNaseH2A was degraded by adding either 0.4 μM 5-ph-IAA (BioAcademia, Japan, #30-003) or 0.5 μM dTAG^V^-1 (Tocris Bioscience, 6914). Control cells were left untreated. Following an additional 48 hrs, cells were harvested by centrifugation. For Pu-seq assays with UV exposure, 4 x 10^4^ cells/ml were seeded. After 48 hrs, 0.4 μM 5-ph-IAA was added. After an additional 4 hours, cells were washed with PBS and exposed to 5 J/m^2^ of UV. Media containing 0.4 μM 5-ph-IAA was replaced and cells were grown for 24 hours before harvesting by centrifugation. Genomic DNA was extracted using the Blood & Cell Culture DNA Midi Kit 100/G Genomic-tips (Qiagen). Small-scale alkaline treatment experiments were conducted by mixing 3 μg genomic DNA with 0.3 M NaOH and incubating the mixture for 2 hrs at 55 °C. Reactions were neutralised by adding 0.3 M Tris-HCl (pH 7.5) and the distribution of ssDNA was analysed using TapeStation RNA ScreenTape (Agilent).

For library preparation, 25 μg of genomic DNA was subjected to alkaline treatment (0.3 M NaOH for 2 hrs) and subsequently separated on a 1.5% agarose gel for 1 hr 40 mins at 100 V. The gel was incubated with 5 μg/mL acridine orange (Invitrogen) for 2 hrs at room temperature with gentle shaking to stain DNA. Destaining was performed overnight in water. DNA fragments ranging from 300-2,000 bp were excised from the gel and purified using the NucleoSpin Gel and PCR Clean-up kit (Macherey-Nagel). Library preparation followed the protocol described previously^21, 22^. The libraries were sequenced as 150-bp paired-end (PE) reads on an Illumina Hiseq X platform (Macrogen, Tokyo, Japan) to generate ∼200 million reads per sample. Two biological replicates were acquired for each condition, except for Pol η-KO and Pu-seq in RPE1 cells, which were conducted once.

### Analysis of polymerase usage

Illumina sequencing reads from Pu-seq data were processed as previously described^19^. Briefly, reads were aligned to the GRCh38 genome with Bowtie2 (version 2.3.5), with reads aligning to multiple locations removed. The 5′ ends of the R1 reads, corresponding to NTP locations, were counted for each 1 kb bin of the genome to provide bin counts for both the Watson and Crick strands. These counts were normalised to the total number of reads for each strand. Pol η-F18A data were then expressed as a ratio of reads in each bin compared to a control sample by dividing each bin count by corresponding count for wild-type Pol η. The resulting data were smoothed using a moving average of the 2 *m* + 1 bins, where *m* is the moving average parameter when specified for individual analyses.

Strand bias scores were calculated as (Pu^W^-Pu^C^)/(Pu^W^+Pu^C^), where R^W^ and R^C^ are the normalised ribonucleotide counts on the Watson and Crick strands, respectively (**Fig. 2a**). Replication asymmetry scores (R.A.S.) at initiation zones were calculated within a 200 kb window on either side of the initiation index peak as (ΣPu^W,R^+ΣPu^C,L^)-(ΣPu^W,L^+ΣPu^C,R^), where ΣPu^W,L^ denotes the sum of Watson strand scores on the left flank and ΣPu^W,R^ the right flank, etc (**Fig.2b**).

### Analysis of cancer mutations

Cancer transcriptome and genome mutation data dataset was acquired from TCGA. R.A.S. scores were calculated as described above, using binned mutation counts in place of ribonucleotide counts. Late replication timing ratio was calculated for each mutation type were calculated as the number of mutations occurring within replication timing bins 12-16 divided by the total number of mutations for that type.

## Acknowledgements

This work was supported by: JST PRESTO grant and FOREST program (JPMJPR18K7 and JPMJFR204X) to YD; JSPS KAKENHI grants (JP21K19203 and JP23H02463) to YD; (JP24K18094) to L.J.B; Naito foundation, Takeda Science Foundation, the Vehicle Racing Commemorative Foundation and Yamada Science Foundation to YD.

## Author contributions

Y.D. conceived the study. L.J.B. and Y.D. designed the experimental approaches. Y.D and L.J.B. designed informatics approaches. L.J.B., Y.M., T.M, M.T. and performed experiments and their analysis. L.J.B. and Y.D. performed computational analysis. L.J.B. and Y.D wrote the manuscript. L.J.B., Y.M. and C.M. edited the manuscript

## Materials & Correspondence

Should be addressed to Y.D.

