## Supplemental figures for "Strand- and replication timing-dependent functions of DNA polymerase η in human DNA replication and mutagenesis"

### Supplementary Figure 1

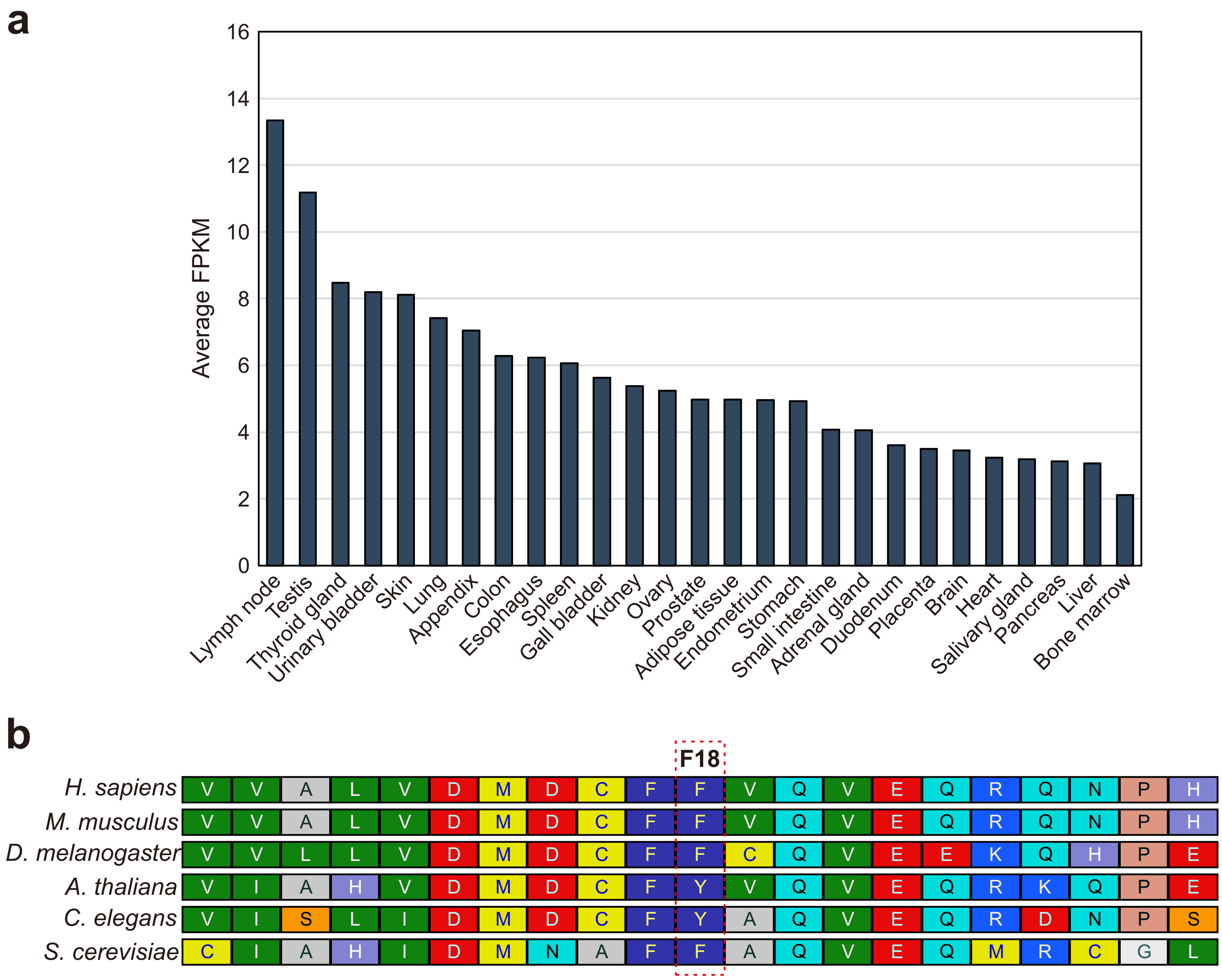


**Supplementary Figure 1 - Analysis of *POLH* expression and sequence conservation. a** Relative abundance of *POLH* mRNA in various human tissues, determined using RNA-seq data from Fagerberg et al. (2014). Expression levels are represented as the average Fragments Per Kilobase of transcript per Million mapped (FPKM). **b** Multiple sequence alignment of the Pol η steric gate region. Conserved proteins were identified using a PSI-BLAST search of the Model Organisms (landmark) database with the human Pol η sequence as the query. Multiple sequence alignment was performed using COBALT. The alignment is colour-coded according to amino acid properties using RasMol colours. Sequences utilised from the NCBI database: *Homo sapiens* (NP_006493.1), *Mus musculus* (NP_109640.1), *Drosophila melanogaster* (NP_649371.2), *Arabidopsis thaliana* (NP_568638.3), *Caenorhabditis elegans* (NP_497480.2) *and Saccharomyces cerevisiae* (NP_010707.3). Database accessed 08/11/2024.

### Supplementary Figure 2

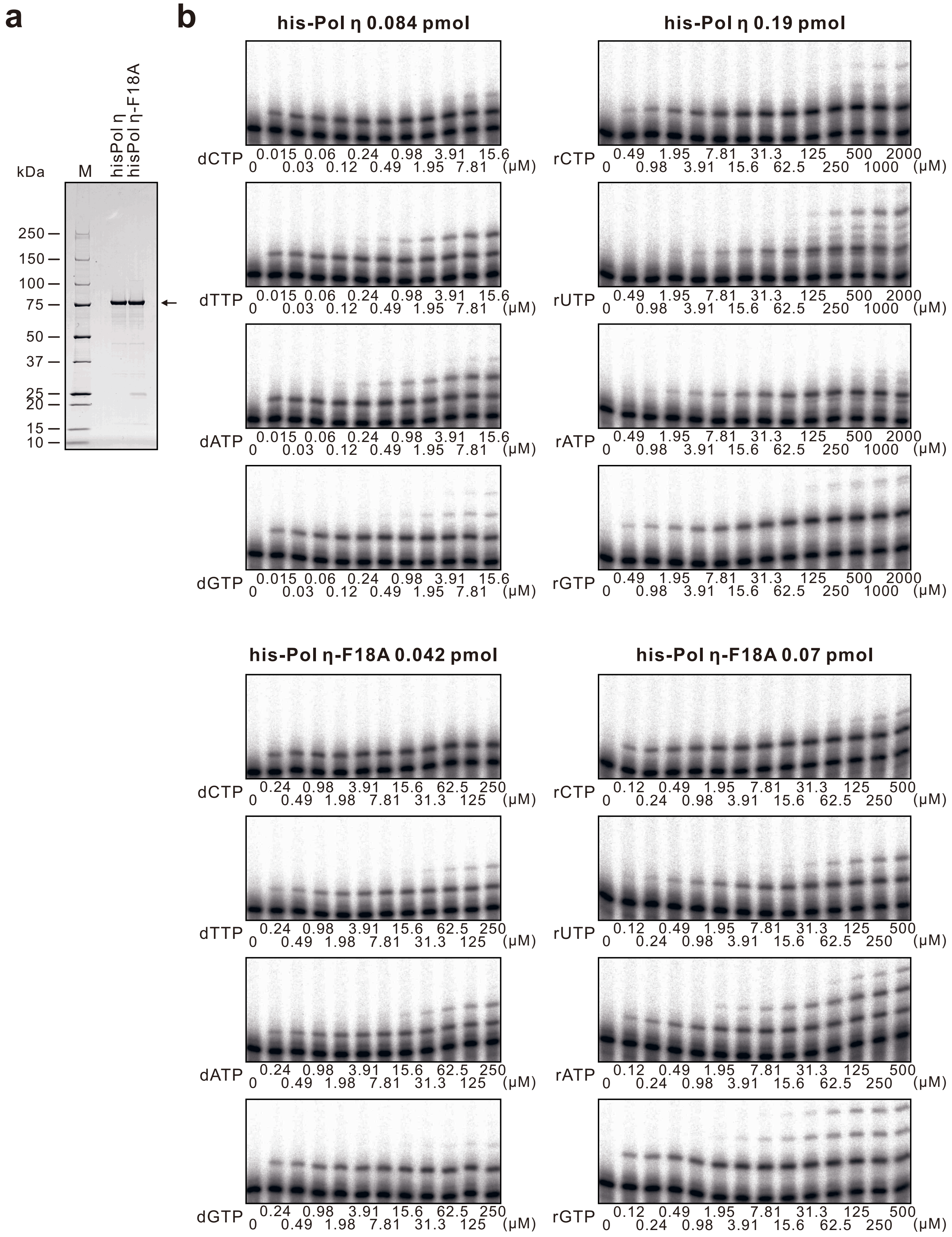

**Supplementary Figure 2 - Single nucleotide incorporation kinetics of Pol η-F18A. a** SDS-PAGE of recombinant His-tagged Pol η and Pol η-F18A expressed and purified from *E. coli*. The expected molecular weight of his-Pol η is ~81 kDa (arrow). **b** Primer extension assays used to calculate single nucleotide incorporation kinetics. A γ-$^{32}$P-labelled 13-mer primer was annealed to 30-mer DNA templates differing at the first templating base was extended by His-tagged Pol η or Pol η-F18A in the presence of the corresponding paired nucleotide and $MgCl_2$.

### Supplementary Figure 3

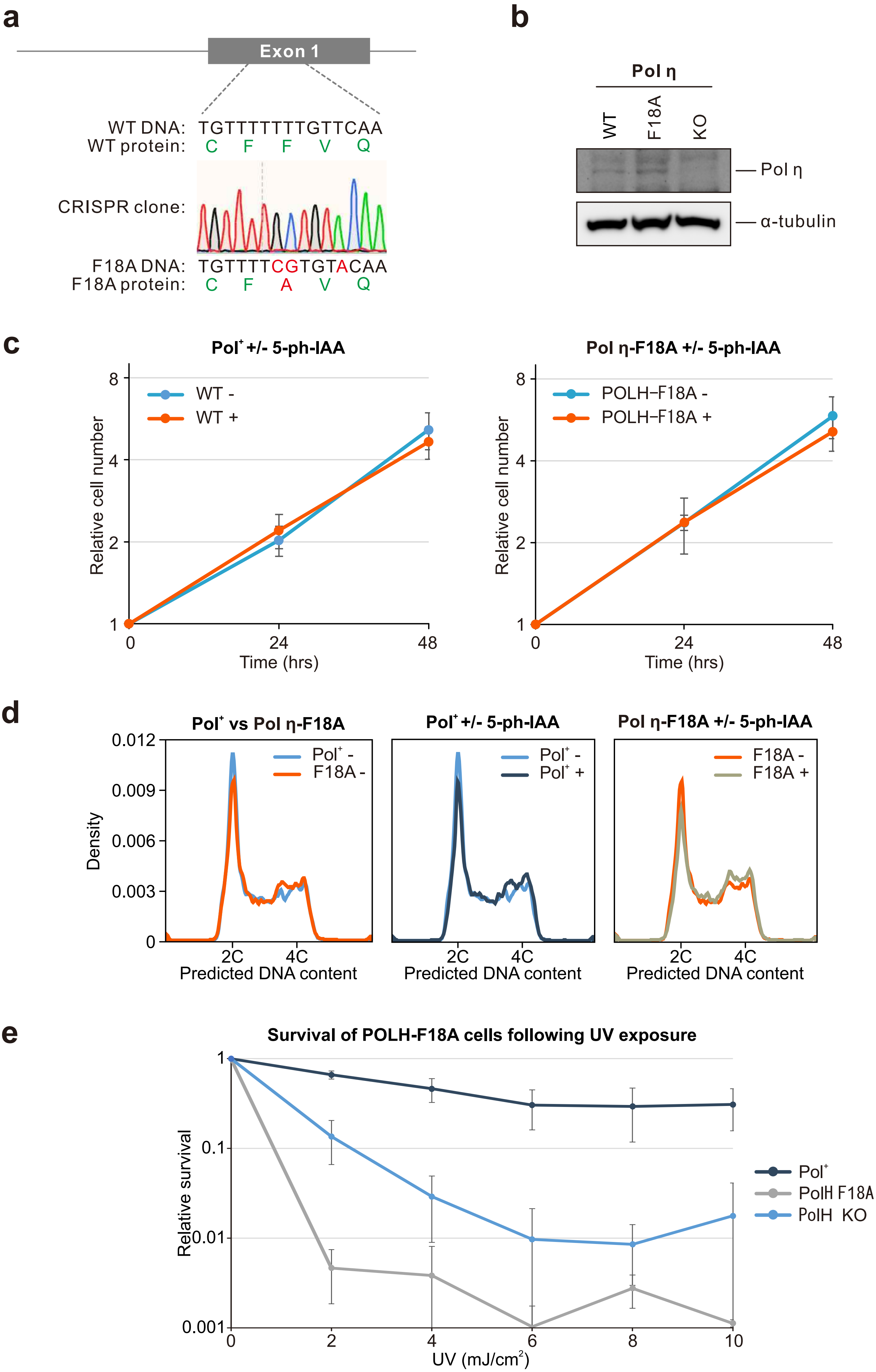

**Supplementary Figure 3 - Characteristics of cells expressing Pol η-F18A. a** Sequencing confirming biallelic POLH-F18A cells generated with CRISPR-Cas12a. **b** Western blot comparing Pol η expression levels in parental Pol η-F18A or Pol η-KO Pu-seq cells. **c** Relative growth of Pu-seq cells in the presence (+) or absence (-) 5-ph-IAA, which induces RNaseH2A degradation. Data represents the mean ± standard deviation from *n* = 3 independent experiments. **d** Cell cycle profile of Pu-seq cells in the presence (+) or absence (-) 5-ph-IAA. **e** Relative survival of Pol η mutant cells exposed to the indicated doses of UV radiation. Survival is expressed relative to untreated controls.

### Supplementary Figure 4

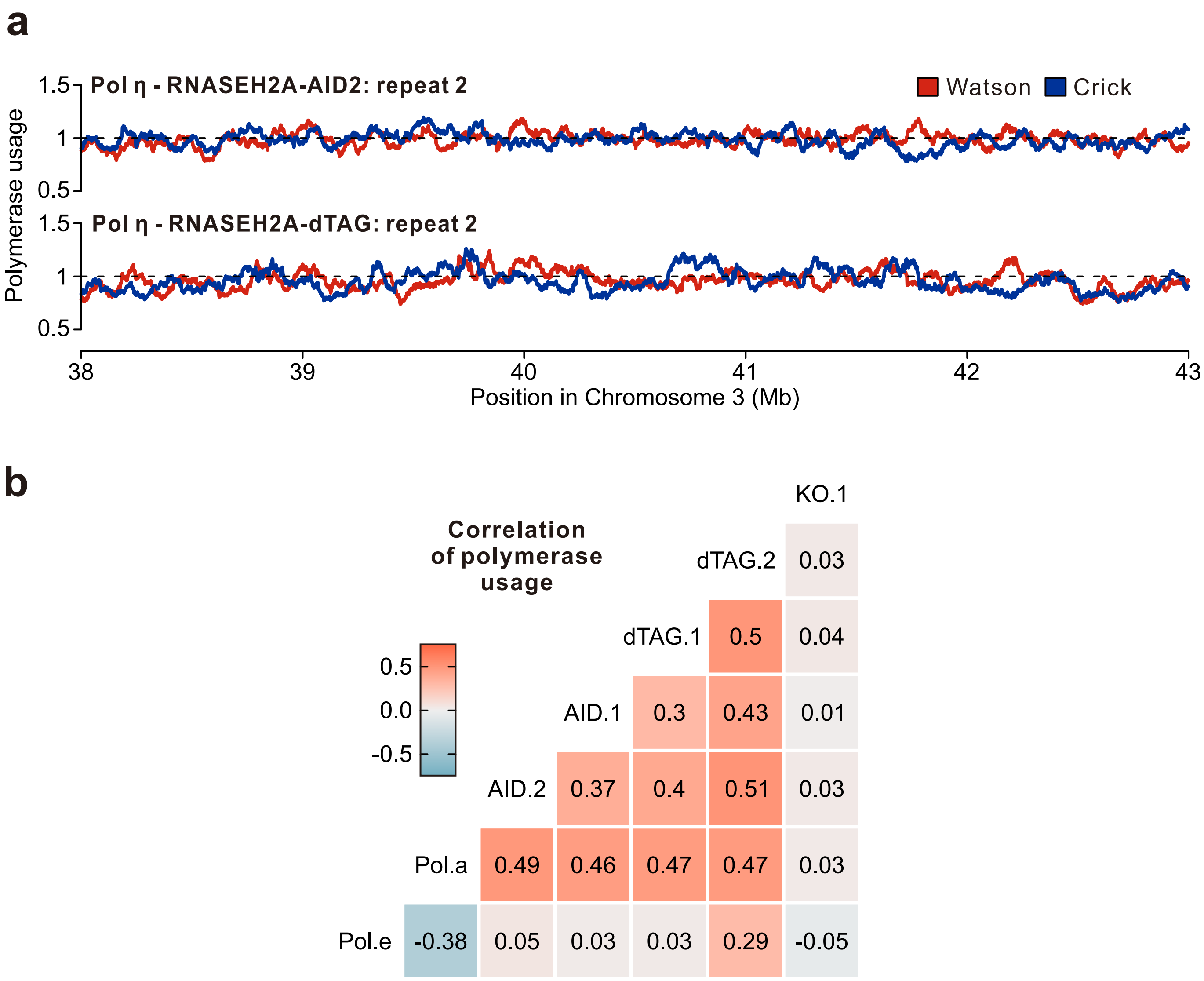


**Supplementary Figure 4 - Comparison of polymerase usage scores in Pol η Pu-seq replicates. a** Smoothed (*m* = 30), strand-specific polymerase usage of Pol η replicates across a representative region of chromosome 3. Red: Watson strand. Blue: Crick strand. **b** Heatmap of Pearson correlations between genome-wide, strand-specific profiles of Pol ε, Pol α and Pol η variants. Red: positive. Blue: negative

### Supplementary Figure 5

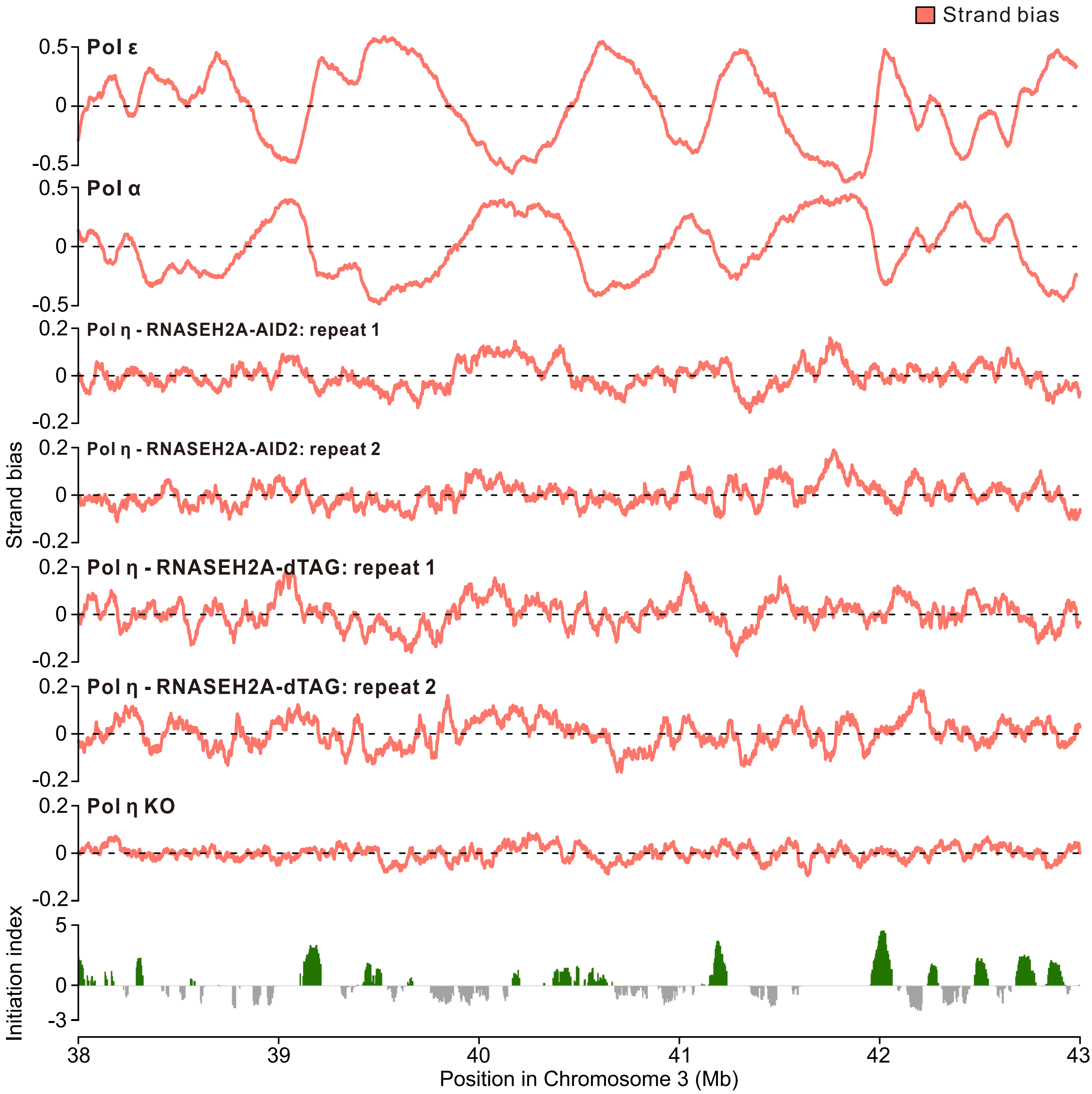


**Supplementary Figure 5 - Strand bias of polymerase usage a region of Chromosome 3.** Smoothed (*m* = 30) strand bias of for Pol ε, Pol α and Pol η usage across a representative region of Chromosome 3. Initiation index (Koyanagi et al., 2022) is also displayed.

### Supplementary Figure 6

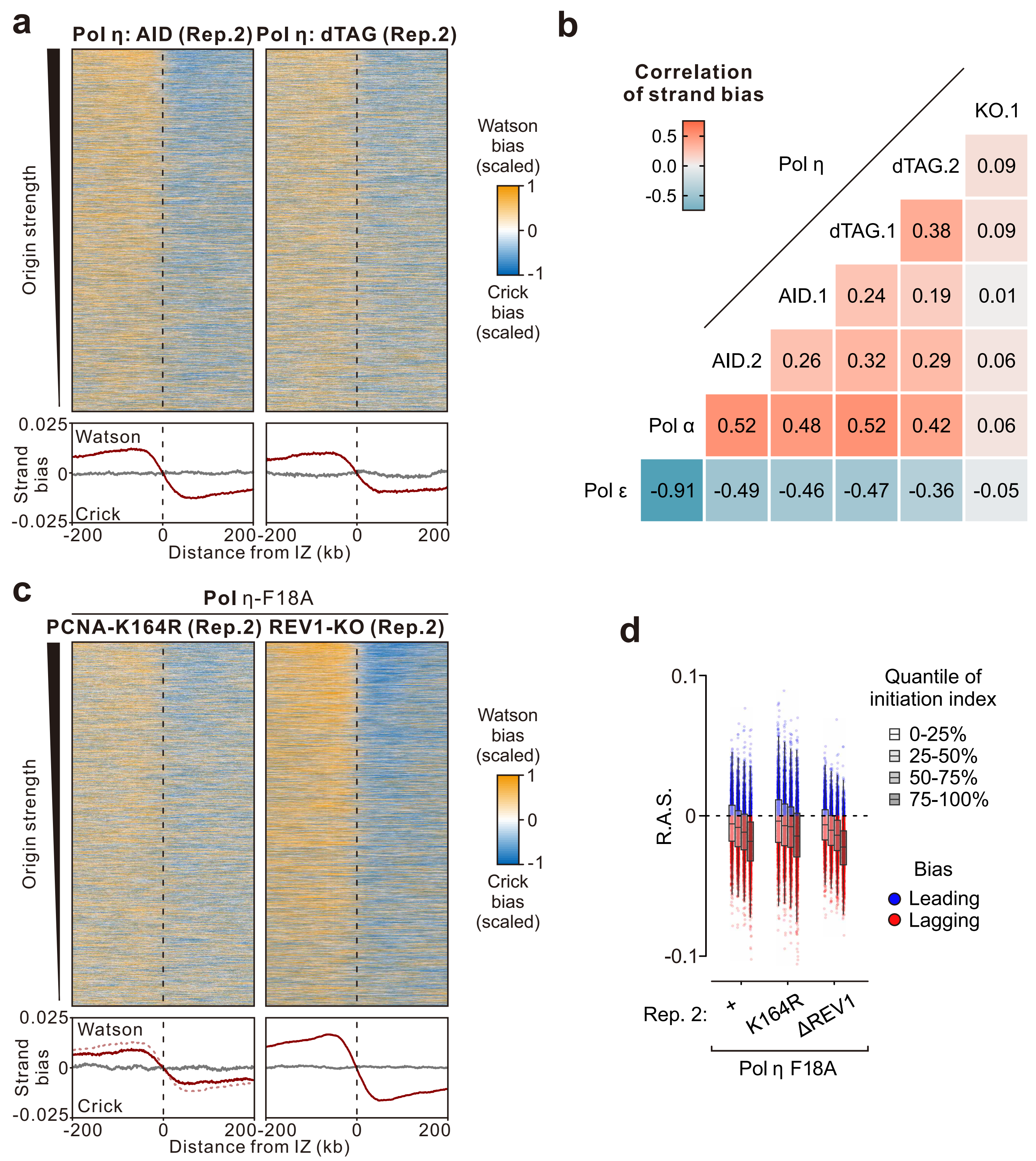


**Supplementary Figure 6 - Strand bias around initiation zones in Pol η Pu-seq replicates. a** Top: Heatmap representation of strand bias scores for Pol η usage ±200 kb around initiation zones (n = 7,139) in replicate samples. The scaled strand bias was ranked by efficiency of replication initiation (top to bottom). Bottom: mean of strand bias around all initiation zones. **b** Heatmap of Pearson correlations between genome-wide strand bias profiles of Pol ε, Pol α and Pol η variants. Red: positive. Blue: negative. **c** As in **a**, but for PCNA-K164R and REV1 KO replicates. **d** R.A.S. of Pol η signal at individual IZ (n = 7,139), quantiled by IZ strength, in replicate samples.

### Supplementary Figure 7

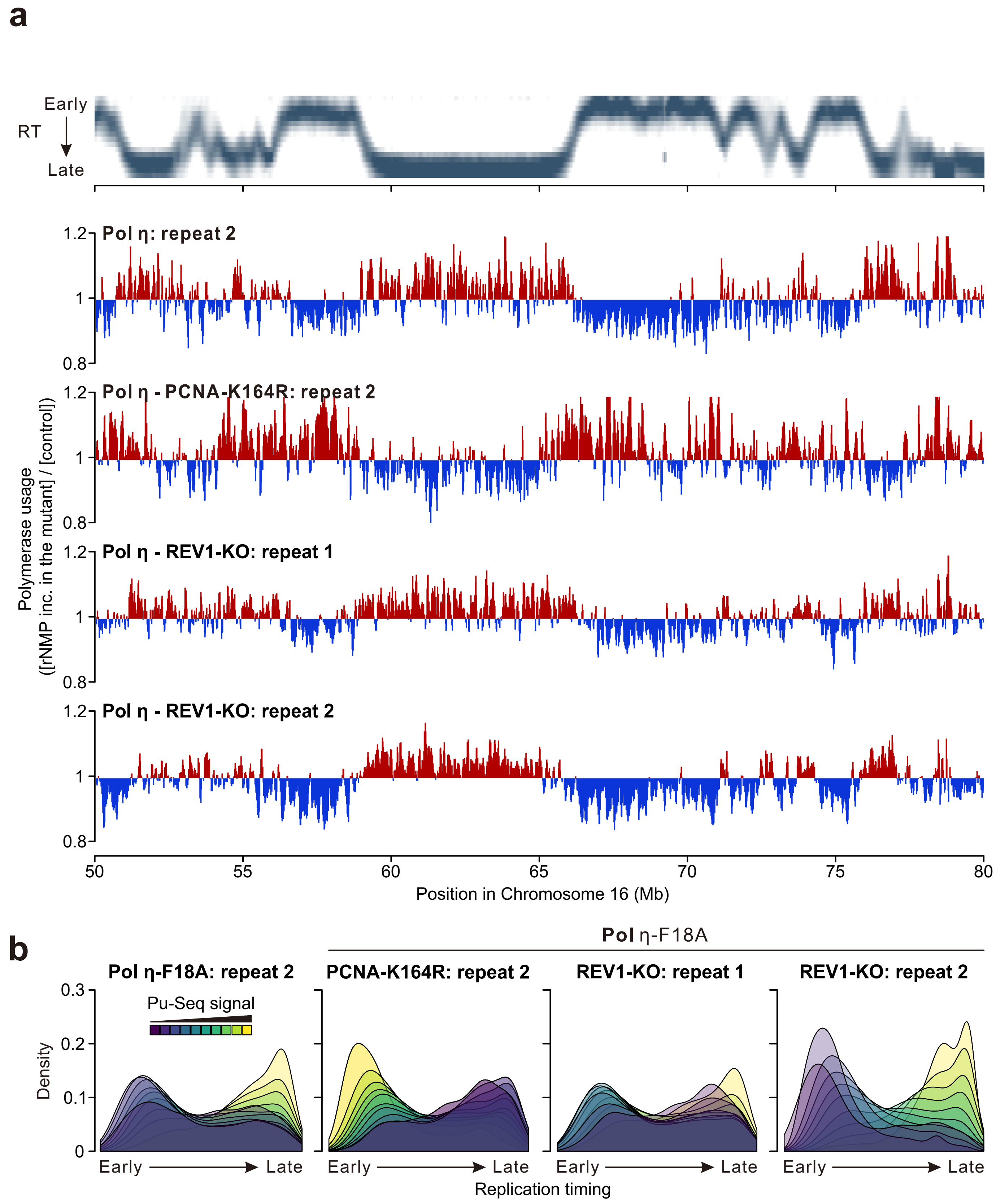


**Supplementary Figure 7 - Pol η usage across replication timing in replicate samples. a** Pol η usage across a representative region of Chromosome 16. Smoothed (m = 30), non-strand-specific Pu-seq profiles show enriched regions in red and depleted in blue. Replication timing (Zhao et al., 2020) data is shown. Data visualised in IGV. **b** Density plots showing replication timing distribution of genomic bins stratified by Pol η Pu-seq signal intensity.

### Supplementary Figure 8

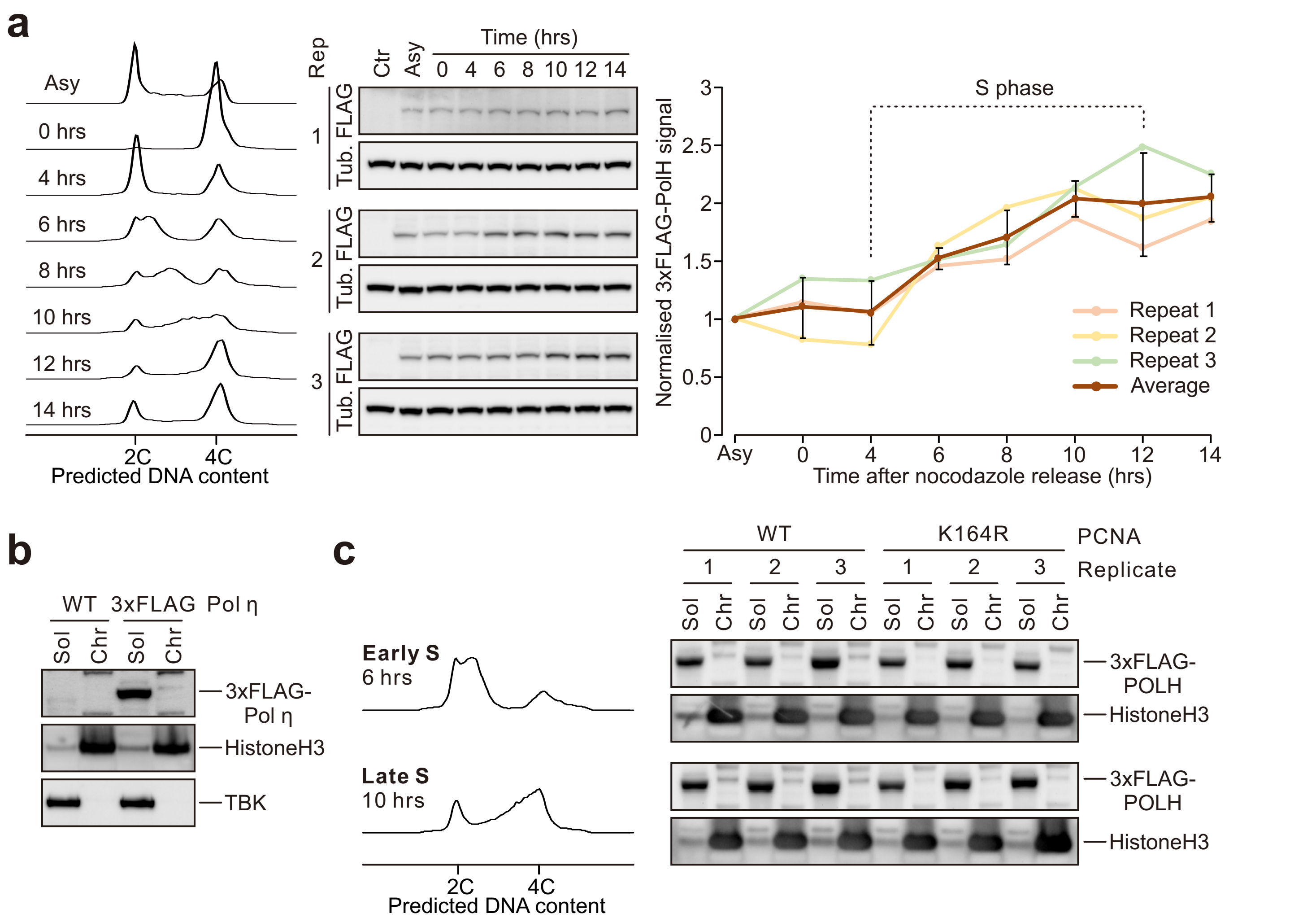


**Supplementary Figure 8 - Expression and chromatin binding of Pol η across S-phase. a** Left: FACS analysis of cell cycle profiles at the indicated time points following release from nocodazole-induced arrest. Middle: Western blots from three biological replicates showing FLAG-tagged Pol η expression from the endogenous locus. Right: Quantification of Pol η levels normalised to α-tubulin and then to the asynchronous (Asy) sample. **b** Western blot detecting chromatin-bound 3xFLAG-Pol η. **c** Chromatin-bound Pol η at early and late stages of S phase following nocodazole release, as confirmed by FACS. Three biological replicates are shown.

### Supplementary Figure 9

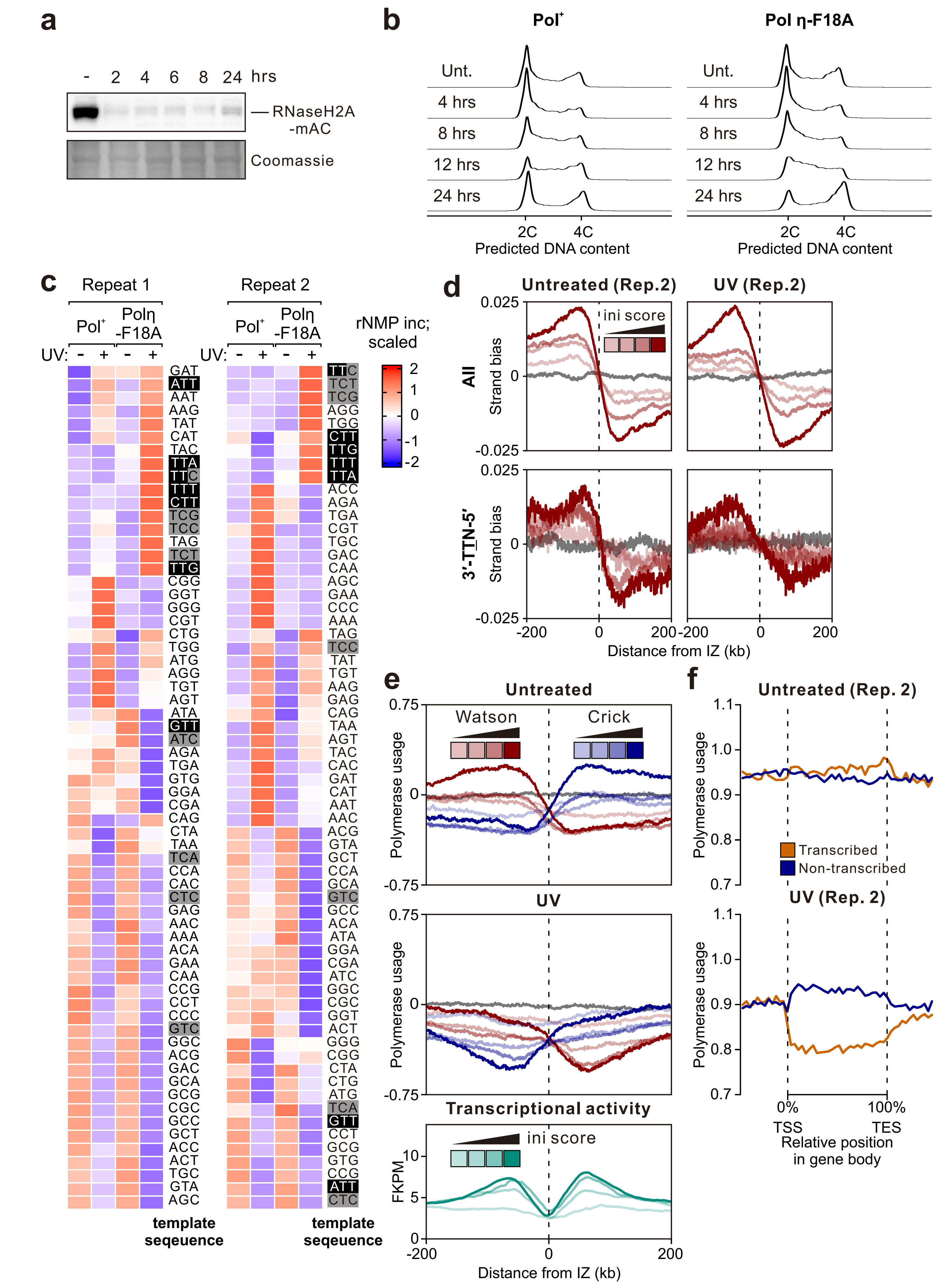

**Supplementary Figure 9 - Usage of Pol η after UV exposure.** **a** Western blot of RNaseH2A-mAC degradation at the indicated timepoints after 5-ph-IAA addition. **b** FACS analysis of cell cycle profiles at various timepoints after exposure to 5 J/$m^2$ of UV radiation. **c** The relative frequency of ribonucleotides detected at each trinucleotide sequence contexts, with and without 5 J/$m^2$ UV, in Pol η-F18A cells and control ($Pol^+$) cells. Frequency of each trinucleotide template within each dataset was calculated, then z normalised by row. Template sequences are displayed 3′ to 5′. **d** Aggregate smoothed (*m* = 30) strand bias of Pol η replicates around all initiation zones (n = 7,139), with and without UV. Data are displayed for all detected ribonucleotides (top) and those deriving from the top TT template sequence motifs (3′-TTA-5′, 3′-TTC-5′, 3′-TTT-5′, 3′-TTG-5′; bottom). Initiation zones are grouped by initiation strength (opacity). **e** Top: Aggregate smoothed (*m* = 30) polymerase usage at sites of replication initiation, grouped by initiation strength (opacity), in the presence and absence of UV. Bottom: Aggregate smoothed (*m* = 30) transcriptional activity (FKPM from RNA-seq), around the same replication initiation zones. **f** Replicate data showing the strand-specific usage of Pol η along gene bodies, scaled to the same arbitrary length. Transcribed and non-transcribed strand usage is shown for the longest 50% of genes among the top 50% most active genes.

### Supplementary Figure 10

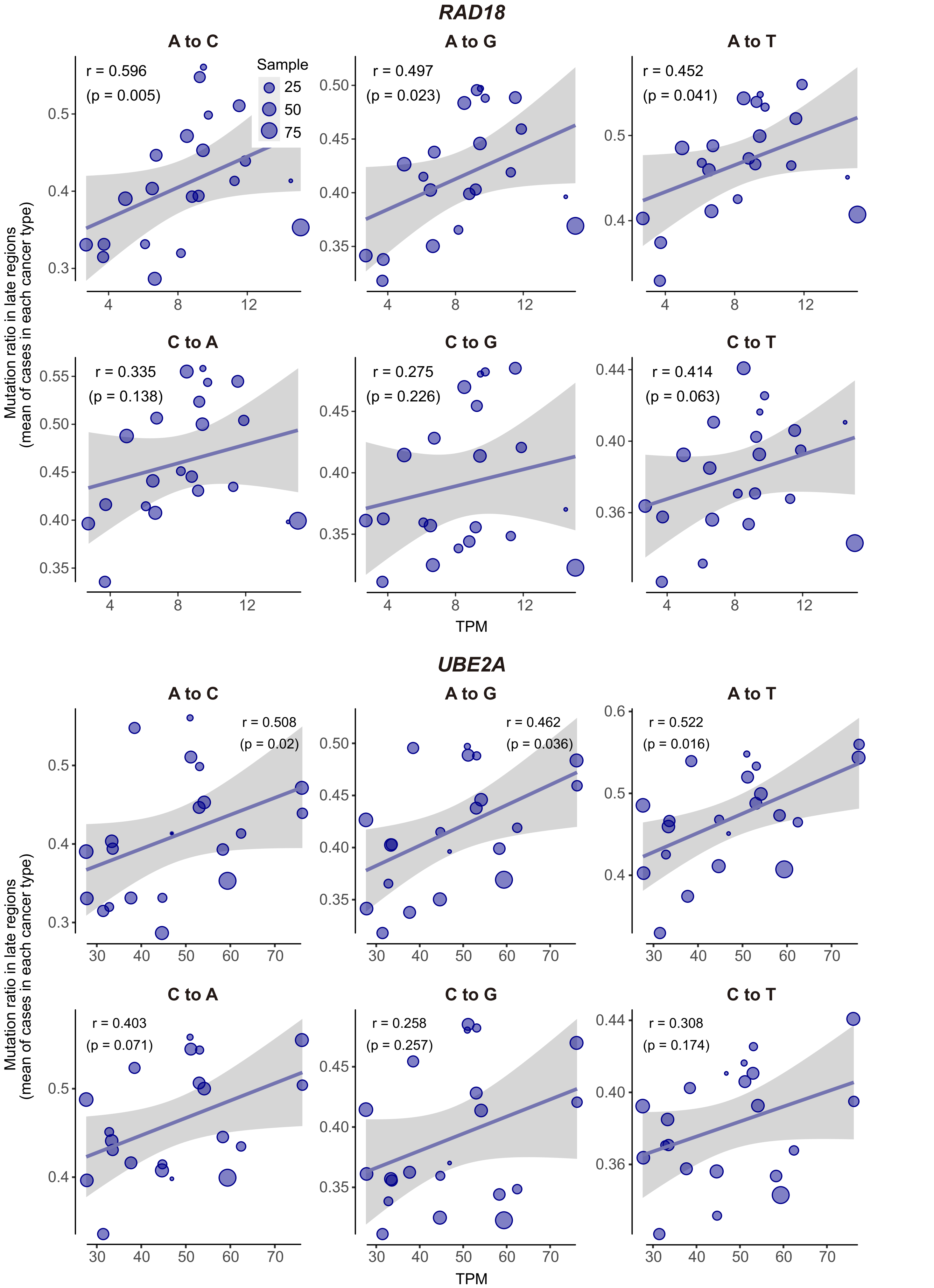


**Supplementary Figure 10 - Correlation between *RAD18* and *UBE2A* expression and mutations in late-replicating regions.** Spearman correlations of *RAD18* and *UBE2A* mRNA expression (TPM) and the ratio of specified mutations occurring in late-replicating domains. Data represent mean values for each cancer type, with point size corresponding to the number of patient samples.
