## Supplemental table 1 for "Strand- and replication timing-dependent functions of DNA polymerase η in human DNA replication and mutagenesis"

Supplementary Table 1

**Resource table**

| **REAGENT** | **SOURCE** | **IDNTFIER** |
| --- | --- | --- |
| **Cell lines** |  |  |
| HCT116 CMV-OsTIR1(F74G)::AAVS1(bi) RNASEH2A-mAID-Clover(bi) | Koyanagi 2022 |  |
| HCT116 CMV-OsTIR1(F74G)::AAVS1(bi) RNASEH2A-mAID-Clover(bi) POLH-F18A(bi) | This study |  |
| HCT116 CMV-OsTIR1(F74G)::AAVS1(bi) RNASEH2A-mAID-Clover(bi) POLH-KO | This study |  |
| HCT116 CMV-OsTIR1(F74G)::AAVS1(bi) RNASEH2A-mAID-Clover(bi) POLH-F18A(bi) PCNA-K164R(bi) | This study |  |
| HCT116 CMV-OsTIR1(F74G)::AAVS1(bi) RNASEH2A-mAID-Clover(bi) POLH-F18A(bi) REV1-KO(bi) | This study |  |
| HCT116 3xFLAG-POLH | This study |  |
| HCT116 3xFLAG-POLH PCNA-K164R(bi) | This study |  |
| RPE1 hTERT EF1-OsTIR1(F74G)::Puro(bi) RNASEH2A-mAID-Clover(bi, Hyg+/Hyg-) | This study |  |
| RPE1 hTERT EF1-OsTIR1(F74G)::Puro(bi) RNASEH2A-mAID-Clover(bi, Hyg+/Hyg-) POLE1-M630F(bi) | This study |  |
| RPE1 hTERT EF1-OsTIR1(F74G)::Puro(bi) RNASEH2A-mAID-Clover(bi, Hyg+/Hyg-) POLA1-Y865F(bi) | This study |  |
| RPE1 hTERT EF1-OsTIR1(F74G)::Puro(bi) RNASEH2A-mAID-Clover(bi, Hyg+/Hyg-) POLH-F18A | This study |  |
| **Oligo DNA** |  |  |
| CACTGACTGTATG | ??? | P13 primer |
| GTGACTGACATACGACTTCTACGACTGCTC | ??? | 30G template |
| GTGACTGACATACAGCTTCTACGACTGCTC | ??? | 30AG template |
| GTGACTGACATACCACTTCTACGACTGCTC | ??? | 30C template |
| GTGACTGACATACTACTTCTACGACTGCTC | ??? | 30T template |
| GTGACTGACATACTACTTCTACGACTGCTC | ??? | 30CPD template |
| GATGAGATTAAGAGCAAGCTTGCCTCCCTGAAGGACGTTCCCAGCCGCATCGAGTGTCCACTCATCTACCACCTGGACGTGGGGGCC**t**T**c**TACCCCAACATCATCCTGACCAACCGCCTGCAGGTGA | IDT Alt-R™ HDR Donor Oligo | ssODN-POLE1-M630F |
| TAGACTTTTTATGACGTGGCTTTTTAATTTCAGGTTTTTATGATAAGTTCATTTTGCTTCTGGACTTCAA**t**AG**c**CTAT**tc**CCTTCCATCATTCAGGAATTTAACATTTGTTTTACAA | IDT Alt-R™ HDR Donor Oligo | ssODN-POLA1-Y865F |
| CCACCCTTCCATGATTTGTACTGTACAACTGCACAAGGTTTATTCCTCAAATGAGGATTTTGCCGCTGCTCCACTTG**t**ACA**gc**AAAACAGTCCATGTCCACGAGAGCAACCACTCGATCC | IDT Alt-R™ HDR Donor Oligo | ssODN-POLH-F18A |
| TACAGCTGTGTAGTAAAGATGCCTTCTGGTGAATTTGCACGTATATGCCGAGATCTCAGCCATATTGGAGATGCTGTTGTAATTTCCTGTGC**gcgc**GACGGAGTGAAATTTTCTGCAAGTGGAGAACT | IDT Alt-R™ HDR Donor Oligo | ssODN-PCNA-K164R |
| CTCAAGCACTTTGGATCCCAGCCATTTCAGATAAGGGATACTCAACCTGTATATTTAAATGCCTATTTCATTCTTGTTAGTTTCCATGCTCCCATGCTCATGGTAACTCATCAGTGAAAGAATGGAATTACAGTTTTCCCTGGCATTTTGGATTAGGTGTTTTTCTAACTGTCCATAAAATGTTGTGTTACAGAAAA**atggactacaaggatcatgatggtgattataaagatcacgacatagactacaaagatgatgacgacaaaggcggaggtggttct**ATGGCTACTGGACAGGATCGAGTGGTTGCTCTCGTGGACATGGACTGTTTTTTTGTTCAAGTGGAGCAGCGGCAAAATCCTCATTTGAGGAATAAACCTTGTGCAGTTGTACAGTACAAATCATGGAAGGGTGGTGGGTATGTATCATTGTTATTGTCACAACTATTCAATGGTAACACAGTGGCACTGTGGCAGGTCCTTTGTT | IDT Alt-R™ HDR Donor Block | HDR-block-POLH-3xFLAG |
| **Oligo RNA** |  |  |
| /AltR1/**CAGGAUGAUGUUGGGGUACA**GUUUUAGAGCUAUGCU/AltR2/ | Alt-R CRISPR-Cas9 crRNA | crRNA-POLE1-M630F |
| /AltR1/UAAUUUCUACUCUUGUAGAU**CUUCUGGACUUCAACAGUCUA**/AltR2/ | Alt-R A.s. Cas12A crRNA | crRNA-POLA1-Y865F |
| /AltR1/UAAUUUCUACUCUUGUAGAU**CCGCUGCUCCACUUGAACAAAAA**/AltR2/ | Alt-R A.s. Cas12A crRNA | crRNA-POLH-F18A |
| /AltR1/**UUUCACUCCGUCUUUUGCAC**GUUUUAGAGCUAUGCU/AltR2/ | Alt-R CRISPR-Cas9 crRNA | crRNA-PCNA-K164R |
| /AltR1/**UGUUACAGAAAAAUGGCUAC**GUUUUAGAGCUAUGCU/AltR2/ | Alt-R CRISPR-Cas9 crRNA  (CD.Cas9.QVNS0030.AQ) | crRNA-POLH-3xFLAG |
| **Plasmid DNA** |  |  |
| pET-h6-POLH | Masuda 2006 |  |
| pET-h6-POLH-F18A | This study |  |
| pSpCas9-POLH (gRNA = GCACAAGTTCGTGAGTCCCG) | This study |  |
| pSpCas9-REV1 (gRNA = GGACACACTCCTAGCTCTAA) | This study |  |
